# PathDiffusion: modeling protein folding pathway using evolution-guided diffusion

**DOI:** 10.64898/2026.01.16.699856

**Authors:** Kailong Zhao, Chenxiao Xiang, Bin Cheng, Yunyun Shen, Wenkai Wang, Shuyun Chen, Baoquan Su, Guijun Zhang, Zhenling Peng, Jianyi Yang

## Abstract

Despite remarkable advances in protein structure prediction, a fundamental question remains unresolved: how do proteins fold from unfolded conformations into their native states? Here, we introduce PathDiffusion, a novel generative framework that simulates protein folding pathways using evolution-guided diffusion models. PathDiffusion first extracts structure-aware evolutionary information from 52 million predicted structures the AlphaFold database. Then an evolution-guided diffusion model with a dual-score fusion strategy is trained to generate high-fidelity folding pathways. Unlike existing deep learning methods, which primarily sample equilibrium ensembles, PathDiffusion explicitly models the temporal evolution of folding. On a benchmark of 52 proteins with experimentally validated folding pathways, PathDiffusion accurately reconstructs the order of folding events. We further demonstrate its versatility across four challenging applications: (1) recapitulating Anton’s molecular dynamics trajectory for 12 fast-folding proteins, (2) reproducing functionally important local folding-unfolding transitions in 20 proteins, (3) characterizing conformational ensembles of 50 intrinsically disordered proteins, and (4) resolving distinct folding mechanisms among 3 TIM-barrel proteins. We anticipate that PathDiffusion will be a valuable tool for probing protein folding mechanisms and dynamics at scale.

## Introduction

Though protein structure prediction has achieved significant advance in recent years, the problem of protein folding remains a grand challenge. Unraveling protein folding pathway is crucial for understanding its function. Although protein function is generally considered to be associated with its native state, increasing evidence suggests that even non-native states also play important biological roles^1, 2^. For example, certain folding intermediates may be precursors to amyloid protein aggregation, which is closely associated with several diseases including Alzheimer’s and Parkinson’s diseases^3, 4^. Therefore, exploring the dynamic process and intermediate states of protein folding is of great importance for understanding disease mechanisms and drug design^5, 6^.

Traditionally, protein folding research has relied on experimental methods such as hydrogen-deuterium exchange, circular dichroism, and fluorescence spectroscopy^7^. Although these techniques provide valuable data on protein dynamics, they often require complex procedures and are limited to a small number of proteins, making it difficult to achieve large-scale investigations in a wider range of protein folding systems^8, 9^. Molecular dynamics (MD) is an important tool for studying folding dynamics. However, it is limited by computational costs, and its time scale is difficult to cover the entire folding process.

In recent years, generative artificial intelligence has significantly advanced the development of the field of protein structure prediction^10, 11^ and protein design^12, 13^. More recently, deep learning models such as BioEmu, AlphaFlow, and DIG have extended these capabilities to generate heterogeneous conformational ensembles^14–17^. Despite these advances, most existing methodologies still prioritize the characterization of thermodynamic equilibrium states over the reconstruction of continuous folding trajectory. We previously developed FoldPAthreader, which explores potential intermediates through folding force field-based conformation sampling^18^. Other emerging methods include generative world models, which learn a strategy from latent spatiotemporal representations to drive folding simulations in latent representations^19^; and the iterative AlphaFold2 strategy, which utilizes implicit energy landscapes to estimate folding pathways based solely on sequence^20^. Despite these methodological advances, developing a generalizable data-driven framework for folding dynamics is still impeded by fundamental constraints, most notably the scarcity of high-resolution experimental data and the computational infeasibility of generating comprehensive training sets via MD^21, 22^.

In this study, we developed PathDiffusion, a novel framework for protein folding pathway prediction using evolution-guided diffusion models. Our method is built on the fundamental relationship between evolution and protein dynamics^18, 23^. Evaluation on 52 proteins with experimental evidence of folding pathway suggests the superior performance of PathDiffusion. Further applications on four datasets related to folding pathway, including fast-folding proteins from Anton’s molecular dynamic simulations, local folding-unfolding transition proteins, intrinsically disordered proteins, and TIM-barrel proteins, confirm that PathDiffusion is promising to explore the landscape protein dynamics.

## Results

### Overview of PathDiffusion

The overall workflow of PathDiffusion is illustrated in **Figure 1**. It comprises two main modules: the first module focuses on preparing position-specific noise schedules (PSNS) from evolutionary information, while the second module iteratively generates the folding pathway using evolution-guided diffusion models. In the first module, we begin with an input protein sequence and construct a multiple sequence alignment (MSA) by searching for homologous sequences in the AFDB50^24, 25^ using MMseqs2^26^ and JackHMMER^27^. Following this, we obtain a multiple structure alignment (MSTA) by aligning the structure models found in AFDB50. Subsequently, PSNS are derived based on the structural consensus and variations observed in the MSTA, which are used to guide the diffusion process in the second module (see Methods for more detail).

**Figure 1.**
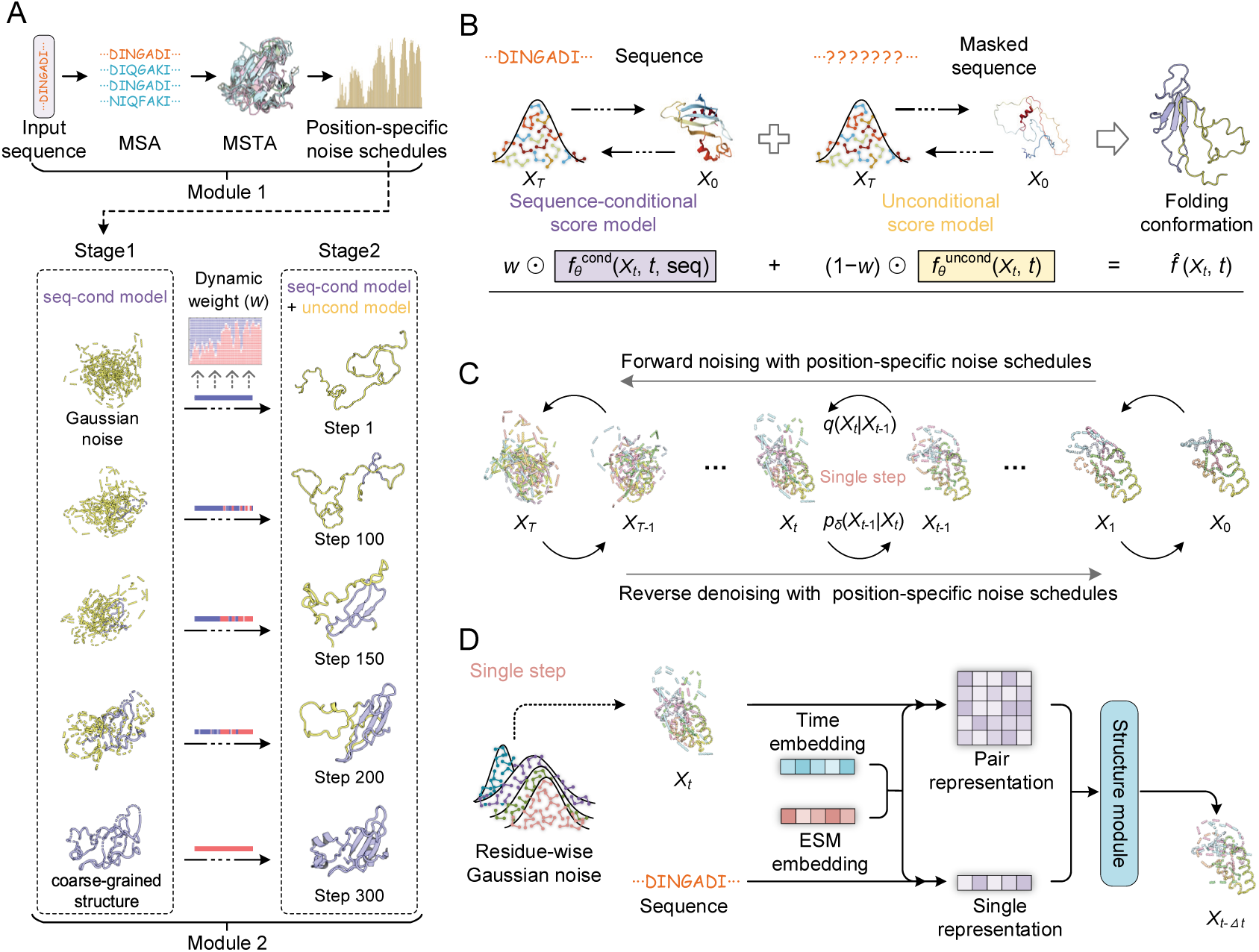
Overview of PathDiffusion. (A) The workflow of PathDiffusion. The framework consists of two modules. Module 1 derives position-specific noise schedules (PSNS) from evolutionary information encoded in multiple sequence alignment (MSA) and multiple structure alignment (MSTA). Module 2 simulates the folding pathway using an evolution-guided, two-stage diffusion strategy. In Stage 1 a sequence-conditional diffusion model is used to generate an initial folding trajectory. In Stage 2, trajectory dynamics are refined via an adaptive dual-score fusion mechanism. (B) Adaptive dual-score fusion mechanism in Stage 2. During denoising, the model combines two score estimators: a sequence-conditional score (directing toward the native folded state) and an unconditional score (directing toward unfolded conformations). These are integrated via a residue-specific adaptive weight ***w*** to produce the final predicted score. (C) Evolution-guided diffusion process. Forward noising and reverse denoising steps are modulated by PSNS, enabling structurally heterogeneous folding rates: evolutionarily conserved regions denoise (structure) earlier and more robustly, while variable regions remain flexible/noisy longer, reflecting biophysical folding principles. (D) Single-step denoising architecture. The model accepts a noisy conformation (and optional sequence) as input. This state, together with timestep embeddings and ESM-derived features, is processed by the structure module to predict the denoised conformation.

The second module employs an evolution-guided two-stage diffusion model to simulate the dynamic process of protein folding, transitioning from unfolded to folded states. In the first stage, a sequence-conditional diffusion model is used to generate an initial folding trajectory. It starts from Gaussian noise and performs *T* steps of evolution-guided reverse denoising. Conserved residues undergo rapid denoising, forming stable local structures in the early steps. Conversely, flexible residues retain noise longer, leading to gradual formation of the global fold as denoising progresses. The resulting *T* denoised states form a continuous trajectory: the initial state (step 1) represents complete Gaussian noise, while the final state (step *T*) corresponds to the fully folded structure. The intermediates (steps 2 to *T*-1) are partially folded, containing varying degrees of noise and local structures, thereby capturing the continuous trajectory of protein folding. In the second stage, an unconditional diffusion model is employed in conjunction with the sequence-conditional model to refine the trajectory dynamics. Adaptive weighting derived from the trajectory in first stage is applied to balance the contributions of both the sequence-conditional and unconditional models. This two-stage pipeline generates continuous trajectory of protein folding, allowing a comprehensive characterization of intermediate states and elucidation of the folding pathway.

### Performance of PathDiffusion

To evaluate PathDiffusion’s capability in modeling protein folding pathways, we compiled a benchmark set consisting of 52 proteins with experimentally characterized folding order from existing literature, termed FP52. Based on experimental evidence, we annotated early folded regions (EFRs) and late folded regions (LFRs) for each protein in FP52, as detailed in **Supplementary Text S1** and **Table S1**.

**Figure 2A** illustrates the dynamic evolution of the protein dihedral angles (*ϕ*, *ψ*) along the PathDiffusion folding trajectory, offering a comprehensive view of how the model explores conformational space and progresses toward the final folded state. Two Ramachandran plots (shown in red) are included as references, depicting the expected dihedral preferences of unfolded and folded states, respectively. The first plot corresponds to unfolded structures and was derived from ∼24,000 structures of intrinsically disordered regions (IDRs) in the human proteome^29^. The second plot represents folded structures, sourced from 20,000 randomly selected structures in PDB. The blue plots show the dihedral angle distributions from PathDiffusion’s folding trajectory, aggregated from conformations at various sampling steps across the 52 benchmark proteins. In the initial stages (steps 1-10), the distributions span a broad range of conformational space, largely consistent with highly unfolded starting conformations. As the denoising process advances, the dihedral angles progressively converge and become enriched in regions characteristic of folded structures, such as β-strands and α-helices, capturing the formation of local secondary structures in intermediate states. At the final stages (steps 290-300), the distributions show strong convergence, closely resembling the patterns observed in PDB structures. This smooth transition in dihedral space demonstrates that PathDiffusion effectively recapitulates the physical folding process, transitioning from a high-entropy disordered state to a low-entropy folded state.

**Figure 2.**
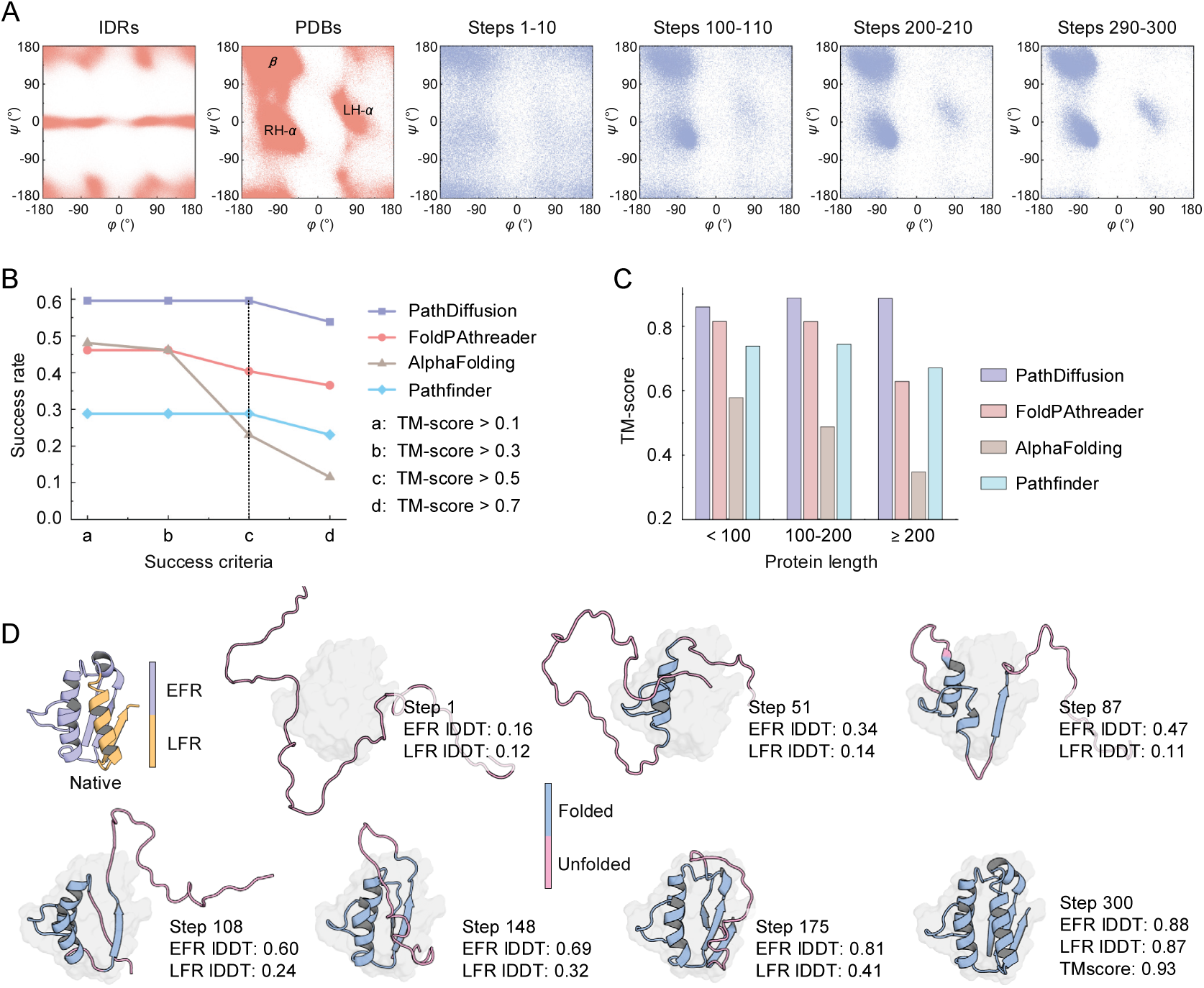
Performance of PathDiffusion on the experimental folding pathway benchmark dataset (FP52). (A) Dynamic evolution of backbone dihedral angles (*ϕ*, *ψ*). Red Ramachandran plots represent reference distributions from intrinsically disordered regions and experimental structures in PDB. Blue plots depict the progressive shift in PathDiffusion-generated conformations, transitioning from a disordered, high-entropy ensemble (early steps 1-10) to a converged, near-native state (late steps 290-300). (B) Comparison of folding pathway prediction success rates on the FP52 benchmark (52 proteins with experimental folding data). (C) Final-state TM-score for proteins with different lengths. (D) Representative folding trajectory generated by PathDiffusion for an example protein (PDB ID: 1AB7_A). lDDT scores computed for early-folding regions (EFRs) and late-folding regions (LFRs) at each step quantitatively demonstrate that evolutionarily conserved EFRs attain stable structure earlier than variable LFRs, consistent with hierarchical folding principles.

We further compare PathDiffusion with several leading methods for folding pathway prediction, including FoldPAthreader^18^, AlphaFolding^20^ and Pathfinder^30^. Both PathDiffusion and FoldPAthreader sample continuous folding trajectory, while AlphaFolding and Pathfinder sample only discrete folding intermediates. To concentrate our evaluation on the critical folding process, we filtered the sampled conformations for all methods. For PathDiffusion, we retained 200 intermediate conformations, omitting the initial 50 and the final 50 steps. For FoldPAtheader, we removed conformations from the initialization and finalization stages, retaining only those from the folding nucleation stage. For AlphaFolding and Pathfinder, we used a clustering strategy with a local Distance Difference Test (lDDT) ^31^ threshold of 0.9 to eliminate redundant structures and select representative conformations.

The performance of the predicted folding pathway is evaluated using success rate. For each conformation, lDDT relative to the native state is calculated separately for EFRs and LFRs (see **Supplementary Text S2**). A conformation is classified as a true positive conformation (TPC) if the lDDT of the EFRs is more than 10% higher than that of the LFRs^21^. A predicted pathway is deemed successful if the TPC ratio (number of TPCs divided by the total conformations) exceeds 0.5 and the TM-score of the final structure surpasses a specified threshold. Finally, the overall success rate is the fraction of proteins in the benchmark dataset with successfully predicted pathways.

Among the 52 test proteins, PathDiffusion consistently achieves the highest success rate across all TM-score thresholds (**Figure 2B**). For instance, at a TM-score threshold of 0.5, the success rates are 59.6%, 40.4%, 23.1% and 28.8% for PathDiffusion, FoldPAthreader, AlphaFolding and Pathfinder, respectively. Detailed results in **Figure 2C** and **Supplementary Figure S2** show that PathDiffusion not only accurately captures the folding order but also produces high-quality structures, yielding the highest TPC ratio (55.7%) and the highest proportion of near-native structures (90.3% with TM-score ≥ 0.7). This hierarchical folding process is further illustrated in **Figure 2D**, which displays the trajectory of a representative protein (PDB ID: 1AB7_A), showing that EFRs attain stable structures (higher lDDT) significantly earlier than LFRs. In contrast, FoldPAthreader has a TPC ratio of 39.3% and a near-native rate of 80.7%. Pathfinder generates a substantial fraction of near-native structures (73.1%) but poorly recapitulated folding order (TPC ratio 29.7%). AlphaFolding, limited in landscape exploration, produces the lowest near-native proportion (21.1%). **Figure 3** illustrates the predicted folding order for 31 proteins successfully modeled by PathDiffusion (TM-score ≥ 0.5), with structures colored by relative folding timing.

**Figure 3.**
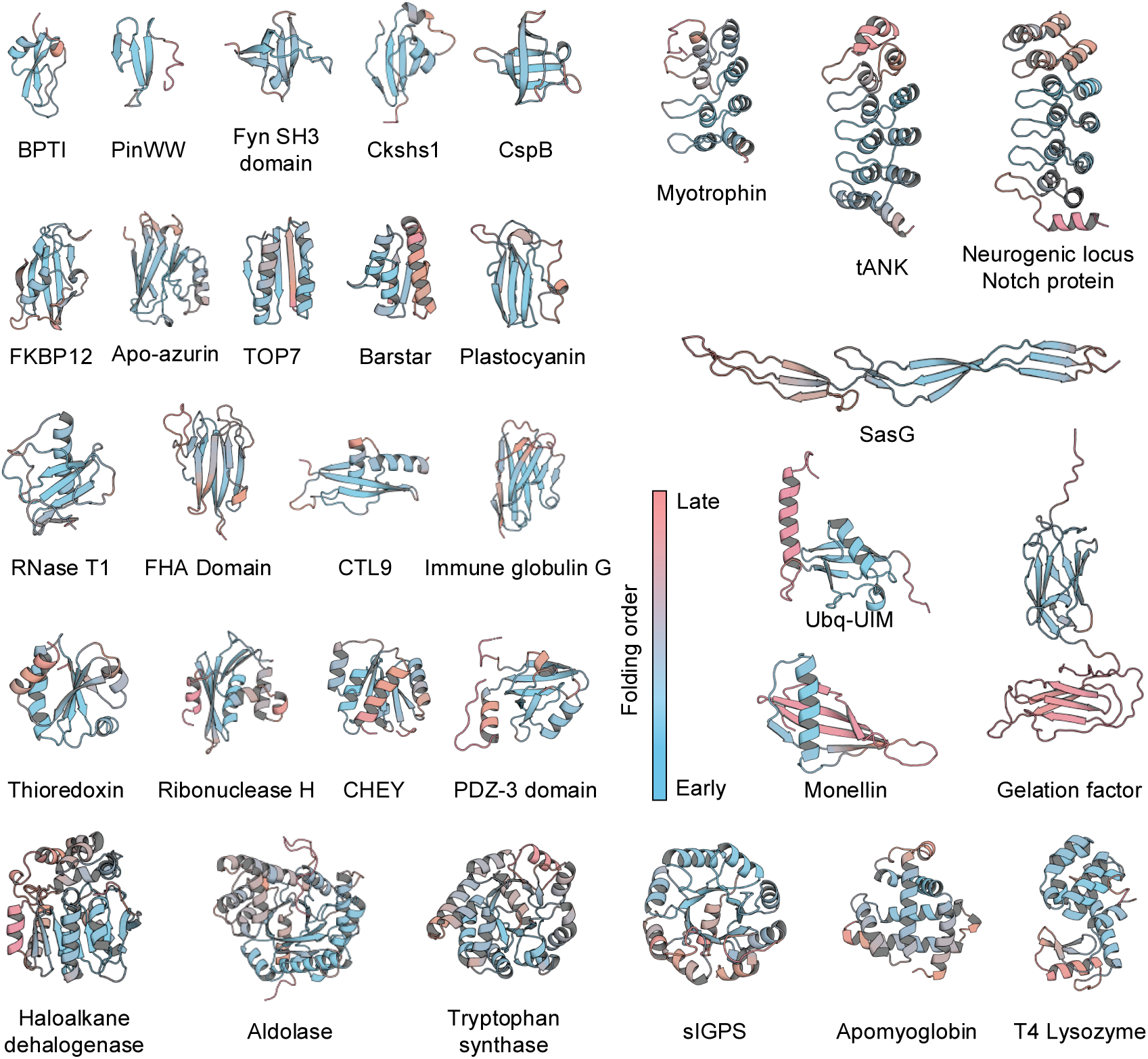
Visualization of the predicted folding order for 31 proteins successfully modeled by PathDiffusion. The structures are colored to illustrate the relative timing derived from the sampled folding trajectory. Blue represents early folding regions, while pink indicates late folding regions. The color gradient is based on the proportion of sampled conformations where the residue aligns within 3 Å of the final state.

### PathDiffusion recapitulates molecular dynamics trajectory generated on the Anton supercomputer

To evaluate PathDiffusion’s ability to capture the equilibrium distribution of folding conformations, we test it on the MD12 dataset, comprising 12 fast-folding proteins that were extensively simulated by MD on the Anton supercomputer^32^. We compare PathDiffusion against BioEmu using its official model, sampling 300 (BioEmu-3×10^2^) and 10,000 (BioEmu-1×10^4^) conformations. For PathDiffusion, we generated continuous folding pathway for each protein comprising 300 conformations, evolving from unfolded to folded states.

To measure the similarity between sampled and reference MD distributions, we calculated the Jensen-Shannon distance (JSD; range 0-1, lower values indicate greater similarity) across three projected feature dimensions: pairwise distance between 𝛼-carbon atoms (PWD); radius of gyration (RG); and time-lagged independent components (TIC), a low-dimensional representation of dynamics^33^.

The quantitative results, presented in **Figure 4A**, demonstrate that PathDiffusion significantly outperforms BioEmu across all projected dimensions. For PWD, PathDiffusion reduces the average JSD by approximately 28.5% and 24.5% compared to BioEmu-3×10^2^ and BioEmu-1×10^4^, respectively. Improvements are more pronounced for RG (62.2% and 59.2% reductions, respectively), which probes structural compactness, and remains strong for TIC (30.5% and 21.9% reductions). The scatter plots in **Figure 4B** further confirm this advantage: for most proteins, PathDiffusion JSD values are above the diagonal, indicating closer alignment with ground-truth MD than BioEmu.

**Figure 4.**
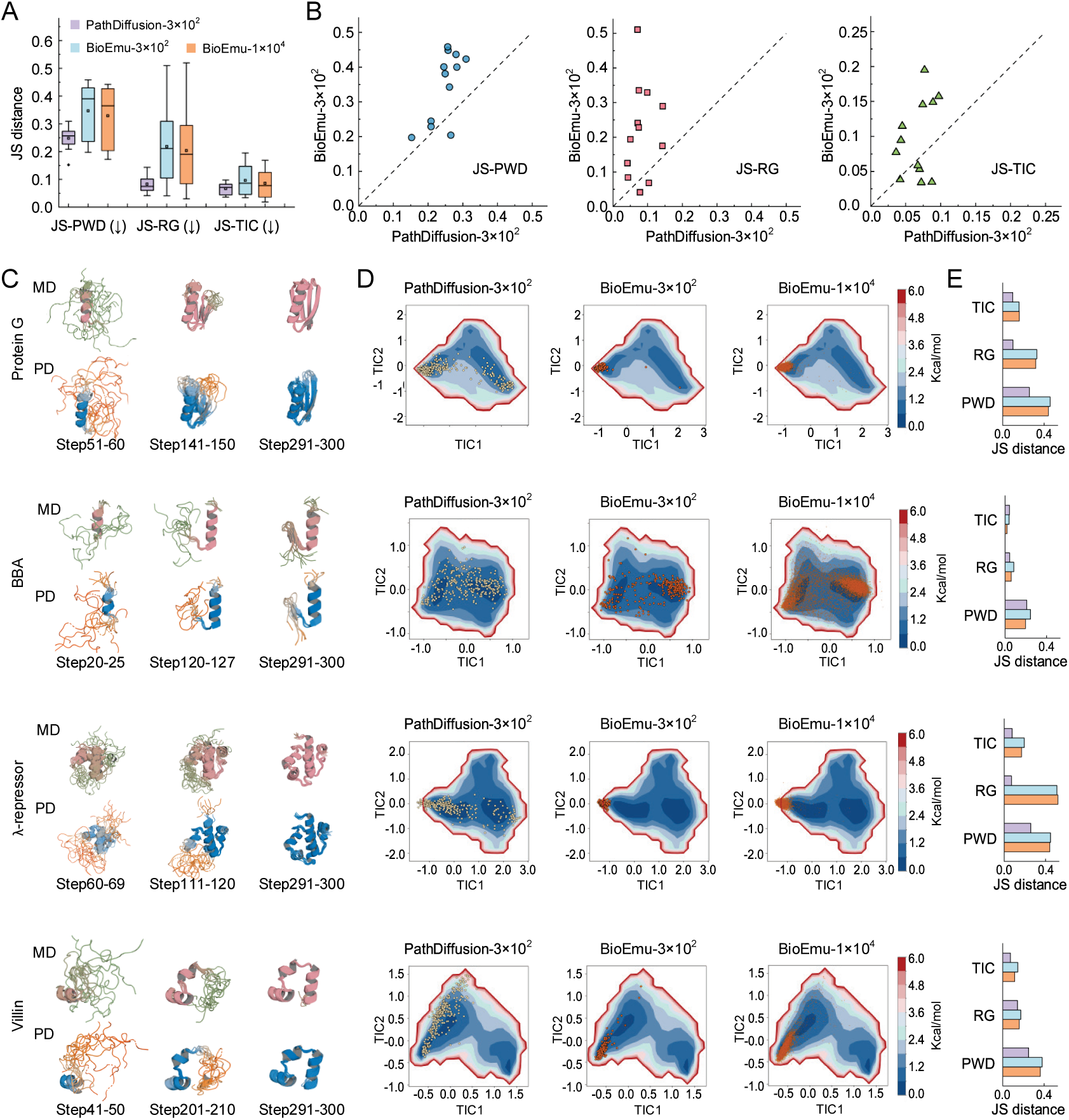
Performance on the molecular dynamics simulation dataset (MD12). (A) Comparison of Jensen-Shannon distances (JSD) between generated ensembles and ground-truth MD simulations across 12 test proteins. A lower JSD value indicates higher similarity to the MD distribution. (B) Head-to-head comparison of JSD for PathDiffusion against BioEmu. Points above the diagonal indicate that PathDiffusion yields distributions closer to the MD ground truth. (C) Visual comparison of MD-simulated ensembles and PathDiffusion folding trajectory at indicated sampling steps. (D) Projection of sampled conformations (scatter points) onto the MD-derived 2D free energy surface. (E) Summary of JSD values for each protein across the three metrics.

To gain deeper insights into the structural distributions, we visualize the 2D free energy surfaces for the 12 proteins (**Figure 4C-E** and **Supplementary Figure S3**). For both *Protein G* and *BBA*, the results reveal a similar folding mechanism, characterized by the preferential formation of helical structures followed by the assembly of β-sheets. In the case of the 𝜆-*repressor*, the first half of helix 1 along with helices 3 and 4 form preferentially, followed by the assembly of the remaining helices. For *Villin*, the process exhibits a sequential formation of helices 1-3.

The free energy surfaces presented in **Figure 4D** illustrate that the conformations generated by PathDiffusion extensively cover the low-energy basins and transition regions, demonstrating a close overlap with the MD data. In contrast, although BioEmu performs comparably for the protein *BBA*, its conformations for other proteins are largely restricted to narrow regions near the native state, which limits its ability to adequately sample the intermediate ensembles. This visual analysis is quantitatively supported by the bar charts in **Figure 4E**, where PathDiffusion yields lower JSD values. This outcome reflects PathDiffusion’s superior capability in capturing the distribution of protein folding conformations compared to BioEmu.

One key advantage of PathDiffusion, beyond accurately estimating conformational distributions, is its ability to generate continuous folding pathways. As illustrated in **Supplementary Figure S4**, mapping the sampling order of PathDiffusion onto the TIC-projected free energy surface reveals that the generated conformations follow the energy gradient, transitioning smoothly from high-energy unfolded states to low-energy stable states. The intermediate structures generated along this pathway closely align with key structures observed in MD simulations. In contrast, BioEmu is restricted to sampling the final equilibrium or metastable states, lacking the capability to provide insights into the folding trajectory or transition states.

### PathDiffusion reproduces local folding-unfolding transitions

The local folding-unfolding transitions of specific regions plays a key role in biological functions, such as the regulation of signaling switches and autoinhibition mechanisms^34, 35^. PathDiffusion explicitly simulates folding dynamics by sequentially sampling the folding pathway, enabling it to capture locally unfolding states. To assess this capability, we compare PathDiffusion with BioEmu^17^, DIG^14^, and AlphaFlow^15^ on the LT20 dataset (derived from the BioEmu study), which includes 20 proteins known to undergo local folding-unfolding transitions. Since local unfolding typically occurs late in folding and/or involves equilibrium fluctuations, we sampled conformations from steps 100-300 in the second stage of PathDiffusion. Evaluations were conducted at two sampling scales: a small batch of 200 conformations and a large batch of 10,000 conformations.

We evaluate the quality of the sampled ensembles using the fraction of native contacts (FNC), as defined in the BioEmu study. For a fair comparison, we only consider samples where the non-local regions remain folded (non-local FNC ≥ 0.5). Following the criteria established by BioEmu, a successful sample was defined as having an FNC below 0.3 for the locally unfolded state and above 0.7 for the folded state. We measured the coverage, defined as the percentage of successful conformations in the top 1% of sampled conformations (ranked by local FNC). With a sample size of 200, PathDiffusion achieves a 72.5% recall rate for accurately identifying locally unfolded states, compared with 65%, 37.5%, and 37.5% for BioEmu, DIG, and AlphaFlow, respectively (**Figure 5A**). While BioEmu has the highest recall for folded states (70%), PathDiffusion remains competitive, correctly identifying 60% of the folded states (**Figure 5B**). Similar trends are observed when the sample size is increased to 10,000, demonstrating that PathDiffusion can efficiently explore the conformational landscape without requiring massive sampling.

**Figure 5.**
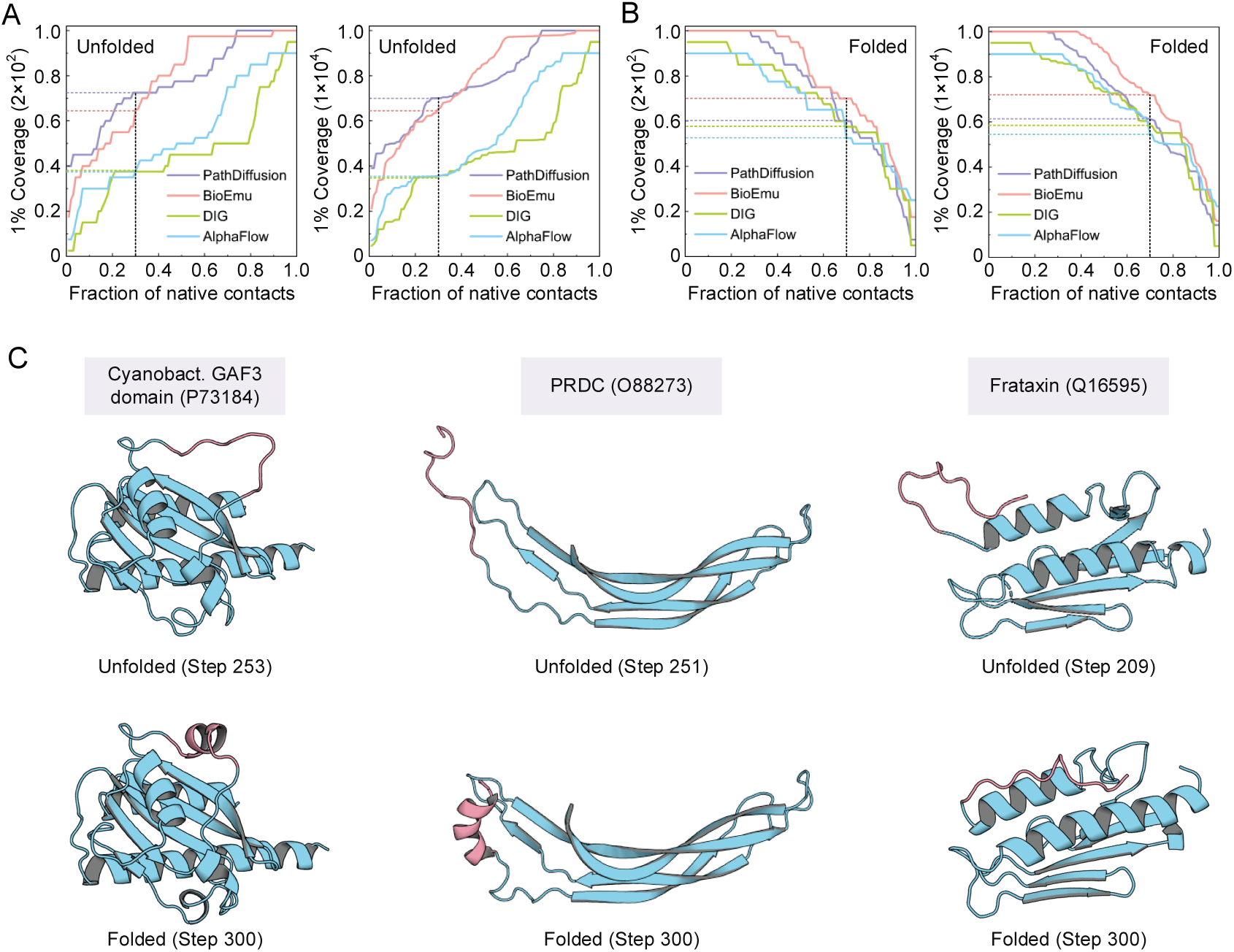
Performance on local folding-unfolding transition dataset (LT20). (A, B) Comparison of the 1% coverage metric at different fraction of native contacts (FNC) thresholds for detecting locally unfolded states and folded states. Evaluations were performed at two sampling scales (2×10^2^ and 1×10^4^). Dashed lines mark the specific FNC thresholds used to define unfolded (< 0.3) and folded (> 0.7) states. (C) Visualization of predicted functional dynamics for representative proteins. Pink segments highlight the specific regions of the native structure known to undergo local unfolding, displayed in both their locally unfolded states and fully folded states.

To validate the biological relevance of PathDiffusion’s predictions, we examine three well-characterized proteins known to undergo local folding-unfolding transitions (**Figure 5C**). The first example is the GAF3 domain of the *cyanobacteriochrome Slr1393* (UniProt ID: P73184), a photoreceptor that regulates red/green light switching. Its photocycle involves major structural rearrangement, in which a long disordered loop in the parental state refolds into a short α-helix in the photoproduct state to stabilize the isomerized chromophore. Crystallographic evidence confirms this disorder-to-order transition^36^. PathDiffusion successfully captures this dynamic feature by generating conformations that represent both flexible loops and folded helices. The second example is the *BMP antagonist PRDC* (also known as Gremlin-2; UniProt ID: O88273). In the ligand-free state, its N-terminal region of PRDC is intrinsically disordered and highly flexible, enabling it to adopt an ordered conformation upon BMP binding that fills the ligand’s binding pocket^37^. PathDiffusion correctly captures this behavior, producing apo-state conformations with pronounced local flexibility in the N-terminus while maintaining a stable, well-structured core. The third example is *human frataxin* (UniProt ID: Q16595), a mitochondrial protein whose dysfunction causes *Friedreich’s ataxia*. Experimental studies have linked pathogenicity to increased flexibility of the C-terminal region, which disrupts mitochondrial iron homeostasis^38^. PathDiffusion sampling consistently reveals elevated local flexibility in this C-terminal segment, in excellent agreement with the experimental observations. Furthermore, visualizations of the predicted results for all 20 proteins are provided in **Supplementary Figure S5**. These cases collectively demonstrate that PathDiffusion not only generates structurally accurate conformations but also reliably identifies functionally relevant flexibility and folding-unfolding transitions.

### PathDiffusion characterizes conformational ensembles of intrinsically disordered proteins

Beyond predicting folding pathways for structured proteins, a critical challenge in protein folding is characterizing the conformational dynamics of intrinsically disordered proteins (IDPs). Because PathDiffusion was trained on both structured and disordered proteins, we applied it to generate ensembles for the dataset IDP50 consisting of 50 IDPs. For comparison, we selected 19 structured proteins from the FP52 dataset with matched sequence lengths as controls.

Analysis of the generated ensembles reveals clear distinctions between structured proteins and IDPs. Root-mean-square fluctuation (RMSF) show that structured proteins rapidly converge to low RMSF values within the first ∼100 steps, stabilizing at levels indicative of rigid, native-like states (**Figure 6A**). In contrast, IDPs sustain markedly higher RMSF throughout the 300-step trajectory, including the final interval (steps 271-300). A parallel pattern emerges in the radius of gyration (Rg, **Figure 6B**), where IDPs exhibit persistently larger Rg values, consistent with their expanded and heterogeneous conformational ensembles.

**Figure 6.**
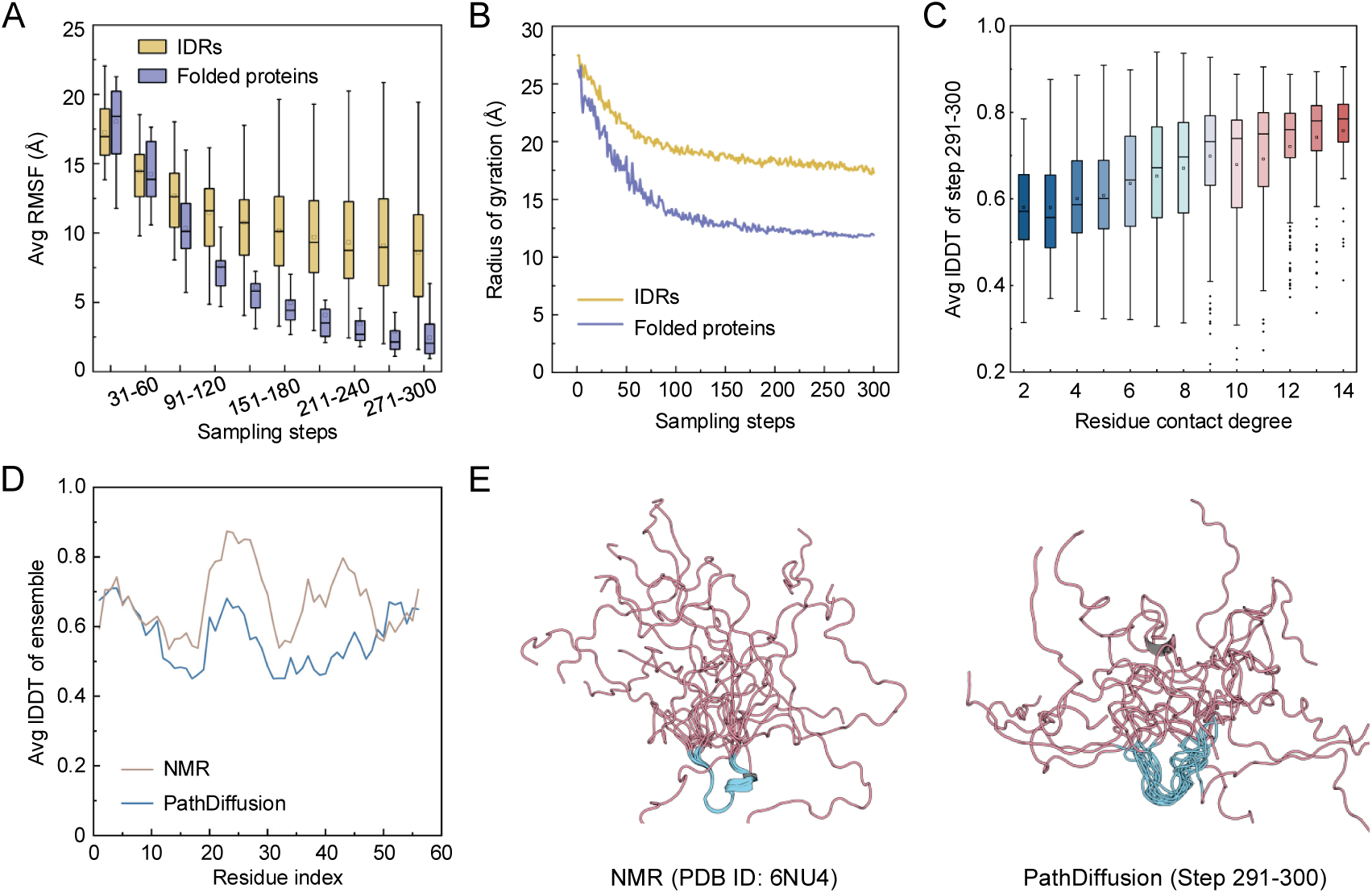
Performance on the intrinsically disordered regions dataset (IDP50). (A) Evolution of average root-mean-square fluctuation (RMSF) calculated in 30-step intervals. (B) Evolution of the radius of gyration (Rg). (C) Box plots showing the relationship between residue contact degree and the average intra-ensemble lDDT (steps 291-300 of IDRs dataset). The contact degree is defined as the number of residues within an 8 Å distance cutoff in the native state. Note that lDDT here quantifies the conformational diversity within the ensemble (lower scores indicate higher flexibility). (D) Comparison of per-residue lDDT for the PathDiffusion ensemble and the reference NMR ensemble. (E) Structural superposition of the NMR ensemble and the PathDiffusion-generated ensemble. Pink represents flexible regions, and blue represents stable regions.

To probe the structural origins of this flexibility, we examined the correlation between per-residue contact degree (based on native contacts) and intra-ensemble lDDT scores (**Figure 6C**). Residues with high contact degrees achieve higher lDDT values, indicating greater structural stability, while those with few contacts show lower lDDT, reflecting elevated flexibility. This demonstrates that PathDiffusion reliably distinguishes locally ordered motifs from fully disordered regions. We conducted an analysis of an example intrinsically disordered protein (IDP) that has experimentally resolved NMR ensembles in PDB (**Figure 6D, E**). It is an *Arabidopsis thaliana RALF8 peptide* (PDB ID: 6NU4), which is primarily disordered but features a short segment that is locally ordered (residues 20-30). PathDiffusion effectively identifies this structured segment while accurately capturing the overall disordered behavior of the peptide. These results highlight PathDiffusion’s robustness in modeling both stable structural elements and flexible regions, facilitating the exploration of complex and heterogeneous conformational landscapes.

### PathDiffusion resolves distinct folding mechanisms in TIM-barrel proteins

Although TIM-barrel proteins are characterized by a highly conserved topology, featuring eight parallel β-strands surrounded by eight α-helices, they exhibit remarkably distinct folding mechanisms. To evaluate whether PathDiffusion can capture these variations in folding pathways, we analyzed three representative TIM-barrel proteins: *indole-3-glycerol phosphate synthase* (sIGPS, PDB ID: 2C3Z)^39^, *α-subunit of tryptophan synthase* (αTS, PDB ID: 1BKS)^40^, and *rabbit muscle aldolase* (Aldolase, PDB ID: 1ADO)^41^. These proteins exhibit high structural similarity (average TM-score 0.63) but low sequence identity (average 0.19), and their folding intermediates have been extensively characterized by hydrogen-deuterium exchange mass spectrometry (HX-MS).

For sIGPS, the folding trajectory simulated by PathDiffusion is highly consistent with the “center-first” folding mechanism revealed by experimental data (**Figure 7A**). HX-MS studies have identified a stable equilibrium intermediate comprising the central regions (βα)_2-5_β_6_, whereas the N- and C-terminal regions remain largely unprotected and disordered^39^. Accordingly, the central segment (residues 50-160) is classified as the EFRs, while the two ends are LFRs. Consistent with this mechanism, PathDiffusion demonstrates a clear hierarchical folding where the lDDT of the EFRs increases significantly faster than that of the LFRs. Structurally, the central β barrel forms a stable core by step 93, whereas the N- and C-termini remain disordered until after step 205. This observation was confirmed by quantitative Z-score analysis (**Supplementary Text S3**), where the PathDiffusion produces high scores (indicating early folding) for the central residues (50-160), almost identical to the high urea protection profile of the native structure.

**Figure 7.**
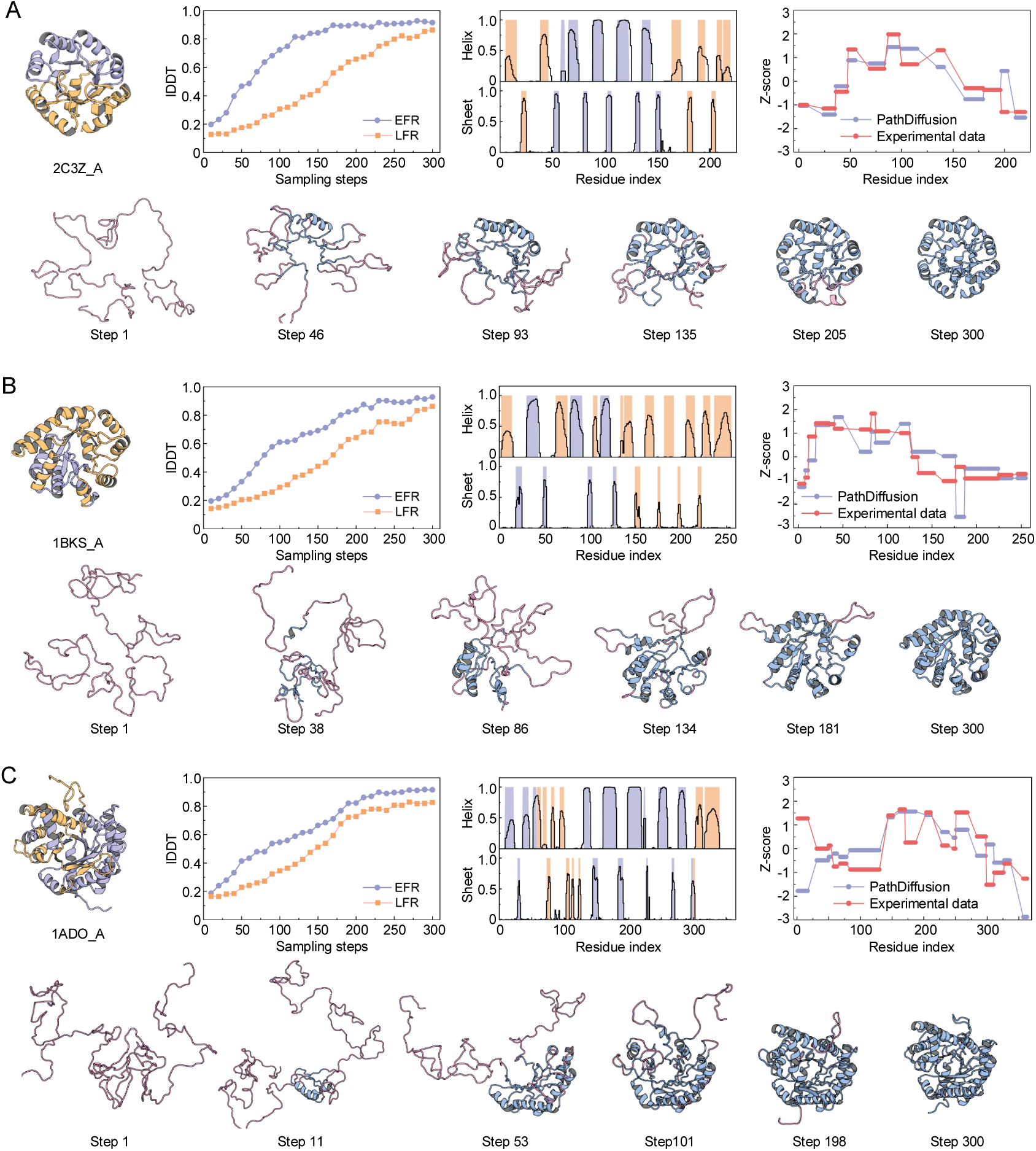
Case studies of folding pathways for three representative TIM-barrel proteins. (A-C) For each protein, the native structure is displayed on the left, colored to distinguish EFRs (blue) and LFRs (orange). The adjacent line plot tracks the lDDT evolution of these regions throughout the sampling process. The bar charts compare predicted secondary structure formation with experimental annotations: the black line indicates the formation frequency predicted by PathDiffusion, while blue and orange background bars denote the native secondary structural elements within experimentally defined EFRs and LFRs, respectively. The Z-score plot compares the predicted folding order with experimental data (see **Supplementary Text S3**). The bottom row presents representative snapshots along the generated folding trajectory, visualizing the continuous transition from a disordered state (pink) to the folded structure (blue).

For αTS, experimental data indicates that the folding intermediate resides in the N-terminal submodule^40^. PathDiffusion recapitulates this this N-terminal preference: lDDT and secondary structure analyses show that the N-terminal domain forms earlier than the C-terminus (**Figure 7B**). Visual inspection of the trajectory reveals that the N-terminal region has formed a stable region at step 86, while the C-terminal region gradually folds in subsequent steps. This asymmetry is further corroborated by the Z-score profile, which shows positive scores at the N-terminus (residues 12-130). Together, these results highlight PathDiffusion’s sensitivity to sequence-specific stability determinants, rather than merely memorizing a generic barrel topology.

For Aldolase, the folding mechanism is notably more complex. Experimental dialysis studies categorize the protein backbone into three stability levels based on their folding rates^41^. Accordingly, we designated regions with the slowest refolding rates (residues 58-130 and 300-363) as LFRs and those with fast or medium rates as EFRs. The PathDiffusion generated trajectory reveals that the EFRs begin to form around step 101, subsequently integrating with other regions (**Figure 7C**). Overall, the predicted folding hierarchy aligns closely with the experimental refolding rates, with one key exception at the N-terminus (residues 1-17). PathDiffusion assigns lower Z-scores (indicating later folding) than observed experimentally. This discrepancy likely stems from quaternary interactions that stabilize the N-terminus in the native homotetramer, which are absent in our monomeric simulations.

In conclusion, although the three TIM-barrel proteins follow different folding pathways, they exhibit common features. All TIM-barrel proteins appear to form a small β-α-β-α module in the early stages. This module provides a basic scaffold, to which other secondary structural elements are added sequentially, ultimately yielding one or more stable intermediates. PathDiffusion successfully detects these early stable regions and predicts their folding order, providing mechanistic insights into how TIM-barrel proteins navigate their complex folding energy landscapes.

### Ablation studies

To quantify the contributions of two key data-driven components (evolution-informed noise schedule and sequence-conditional score model) in PathDiffusion, we conduct ablation studies (**Figure 8A**). The default PathDiffusion achieves a baseline success rate of 59.6% at a TPC threshold of 0.5. When both the evolution-informed noise schedule and the score model are removed, the performance drop to the lowest level, with a success rate of only 11.5%.

**Figure 8.**
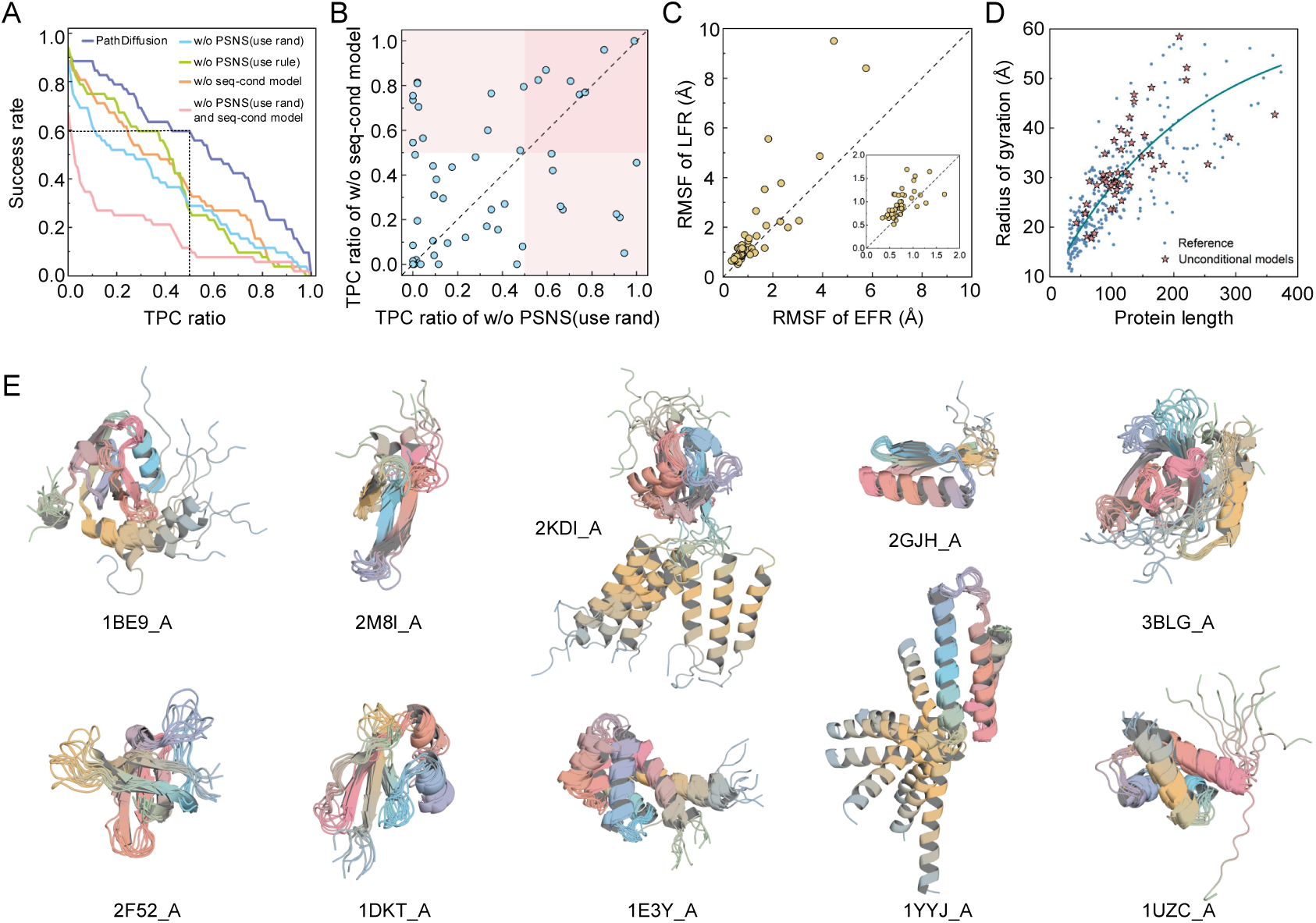
Results of ablation study. (A) Contributions of evolution-informed noise schedule and sequence-conditional score model. The curves display the success rate across varying TPC ratio thresholds. The dashed lines indicate the default performance baseline at an TPC threshold of 0.5. (B) Scatter plot compares the TPC ratio achieved when sampling without using PSNS versus without using a sequential conditional scoring model. The pink shaded region indicates targets where at least one component achieves an TPC ratio greater than 0.5. (C) Conformational diversity analysis of the sequence-conditional model. The scatter plot compares the RMSF of EFRs against LFRs within the generated equilibrium ensembles. Points above the diagonal line indicate that LFRs exhibit higher structural flexibility than EFRs. (D) The relationship between Rg and length of proteins sampled by the unconditional score model. (E) Visualization of representative conformational ensembles generated by the sequence-conditional model. These superimposed structures illustrate the equilibrium fluctuations quantified in (B).

#### Evolution-informed noise schedule

We investigate the impact of the evolution-informed noise schedule (i.e., PSNS). We first replace PSNS by random noise schedule. In this setting, the target sequence is segmented based on predicted secondary structures, and random noise schedules are assigned to the residues within each segment. This leads to a sharp decline in success rate from 59.6% to 28.8%. To test if a general physical prior could bridge this gap, we then implement a rule-based schedule. We introduce a rule-based noise schedule that imposes sequential ordering from N- to C-terminus and a structural hierarchy assigning progressively lower noise to α-helices, β-sheets, and loops, respectively. Although this rule-based schedule outperforms the random schedule at lower TPC thresholds, it yielded only comparable results at higher thresholds, and both strategies falling far short of the full model. This result highlights that general physical rules cannot substitute for the position-specific evolutionary signal encoded in PSNS. The PSNS, derived from multiple structure alignment, is essential for guiding PathDiffusion toward the correct folding pathway of the target protein.

#### Sequence-conditional score model

We next ablate the sequence-conditional score network by replacing its predicted scores with scores computed directly from the static ground-truth structure (see Section 2.3). This change reduces the success rate to 34.6% at a TPC threshold of 0.5, indicating that the learned score model captures evolutionary information about folding dynamics that is superior to static geometric priors derived solely from the native state.

Examination of TPC distributions for models relying on only one component further confirms that that the two components are complementary (**Figure 8B**), thus combining both sources yields markedly superior pathway reconstruction compared to either alone. These ablation experiments demonstrate that PathDiffusion’s strong performance arises from the tight synergy between the evolution-informed noise schedule and the sequence-conditioned score model.

#### Generative capabilities of the score models

We further evaluate the generative capabilities of the sequence-conditional score model and the unconditional score model. Both models are trained to convergence, using 100 denoising steps for structure generation. For the sequence-conditional score model, we utilized a uniform noise schedule (applying the same noise level to all residues) to generate 100 structures for each target. As shown in **Figure 8C**, the average RMSF for EFRs and LFRs were 1.205 Å and 1.676 Å, respectively, with 81% of targets exhibiting higher RMSF in LFRs than in EFRs. This indicates that the sequence-conditional model effectively captures intrinsic structural flexibility, revealing that LFRs typically exhibit greater conformational diversity than EFRs during functional dynamics with representative ensembles shown in **Figure 8E**. For the unconditional score model, we generate one conformation per target and calculated the radius of gyration (Rg) (**Figure 8D**). The resulting Rg values increase with protein length in close agreement with the statistical scaling laws observed for intrinsically disordered proteins^42, 43^. This suggests that the unconditional model has successfully learned the generic physical properties and spatial scaling constraints that govern expanded and disordered conformations.

### Analysis of compactional efficiency

To evaluate the practical utility of PathDiffusion, we benchmarked its computational efficiency. All experiments were conducted locally on an A100 GPU with 40 GB of memory. We measured the runtime required to sample folded trajectory on the FP52 dataset using the default parameters for each method. As illustrated in **Supplementary Figure S6A**, PathDiffusion is slower than AlphaFolding but 5-10 times faster than both FoldPAthreader and Pathfinder.

We further assessed the computational cost of generating extensive conformational ensembles for 12 fast-folding proteins (**Supplementary Figure S6B**). PathDiffusion samples 10,000 conformations in under 3 hours. Although slower than BioEmu (< l.5 hours) due to its more extensive denoising schedule (100 steps versus 30-50 for BioEmu), both methods operate on comparable timescales suitable for practical use. In contrast, previous studies have reported that all-atom MD simulations require 2,000-100,000 GPU hours to produce equivalent ensembles^17^. Consequently, PathDiffusion achieves an acceleration of four to five orders of magnitude compared to traditional MD simulations. In summary, PathDiffusion effectively recovers protein conformational ensembles and elucidates folding pathways with high accuracy, offering substantial potential for investigating protein folding pathway and dynamics.

## Conclusions

In this study, we introduce PathDiffusion, a generative diffusion framework that directly predicts continuous protein folding pathway from amino acid sequence. By incorporating residue-level evolutionary information into a two-stage diffusion process, PathDiffusion efficiently explores protein folding pathway while preserving high native-state accuracy. Unlike existing methods that primarily sample near-native equilibrium distributions, PathDiffusion explicitly simulates the dynamic process of folding. It allocates more sampling resources to exploring unfolded states and transition regions as well as nucleation events, making the method more suitable for studying folding mechanisms that evolve over time.

PathDiffusion does not rely on MD simulations data for training or fine-tuning. It predicts folding pathways in a zero-shot manner, achieving several orders of magnitude speed improvement while maintaining strong agreement with MD-derived conformational ensembles. Compared to existing deep learning-based ensemble generators, PathDiffusion generates a richer variety of intermediates and can distinguish different folding mechanisms even among topologically similar proteins. PathDiffusion can also reproduce function-related local unfolding-folding transition events, which are crucial for allosteric regulation and misfolding-related diseases. Furthermore, extensive evaluations on intrinsically disordered proteins demonstrate that PathDiffusion effectively reproduces the intrinsic structural flexibility and conformational heterogeneity of disordered regions. Collectively, these capabilities position PathDiffusion as a powerful bridge linking sequence information to structural dynamics and biological function.

Despite these advances, limitations remain. The current framework relies on MSA and homologous structures to estimate the evolutionary information, reducing accuracy for orphan proteins or *de novo* designs lacking evolutionary context. Future work could develop single-sequence dynamical inference from predicted features^44^. Additionally, diffusion steps represent abstract progression rather than physical time, precluding direct prediction of absolute folding rates or metastable state lifetimes. Future extensions could incorporate biophysical principles, such as coarse-grained energy estimation or kinetic network models, to map generated pathways onto a physically-based free energy landscape and provide approximate kinetic information^45, 46^.

## Methods

### 1. Training and test data

#### Training datasets

PathDiffusion employs two specialized diffusion models: a sequence-conditional diffusion model for folding structured regions and a conditional diffusion model for handling unstructured (intrinsically disordered) regions. To train these models, we constructed two dedicated datasets. The structured dataset consists of ∼530,000 structures from PDB and the unstructured dataset contains ∼24,000 intrinsically disordered regions from CALVADOS coarse-grained simulations^29^. Detailed procedures for dataset construction, including filtering criteria and processing steps, are provided in the Supplementary Information.

#### Test datasets

PathDiffusion are evaluated on four independent benchmark datasets that cover diverse aspects of protein folding dynamics, which are briefly introduced below. FP52: 52 proteins with experimentally determined folding pathways, curated from the literature; MD12: 12 fast-folding proteins with long-timescale molecular dynamics trajectory simulated on the Anton supercomputer^32^; LT20: 20 proteins exhibiting local folding-unfolding transitions, selected from the BioEmu study^17^; IDP50: 50 intrinsically disordered proteins sourced from PDB. Additional details on dataset composition, selection criteria, and preprocessing are available in the Supplementary Information.

### 2. PathDiffusion algorithm

PathDiffusion works through two major modules (**Figure 1**), where the first module prepares a position-specific noise schedules (PSNS) from evolutionary information and the second module iteratively generate folding pathway using PSNS-guided two-stage diffusion models.

#### 2.1 Derivation of PSNS from evolutionary information

In the first module (**Figure 1A** top panel), we derive the PSNS for a target protein from its evolutionary information captured by multiple sequence alignment (MSA) and multiple structure alignment (MSTA).

##### MSA generation

For a target protein sequence, an MSA is first constructed by searching for homologous sequences in AFDB50, a clustered subset of the AlphaFold Protein Structure Database containing over 52 million representative structures selected at 50% sequence identity. The search is performed using MMseqs2^26^ and JackHMMER^27^, with search hits ranked according to sequence identity and E-value. Specific parameters are set to non-default values to ensure search quality: for MMseqs2 (--evalue 1 --sensitivity 6.0 --sort-results 1 --max_seqs 10000) and for JackHMMER (-N 3 -E 1). In practice, we prioritize homologous sequences identified by MMseqs2; JackHMMER is utilized only if fewer than 10 sequences are returned by MMseqs2.

##### MSTA construction and PSNS derivation

We construct an MSTA by aligning the corresponding structures of the identified homologs from AFDB50. Specifically, the structure of the top-ranked protein in the MSA serves as the reference. The remaining homologs are structurally aligned to this reference using TM-align^28^, resulting in the MSTA. The PSNS is then derived from the MSTA using the following formula:

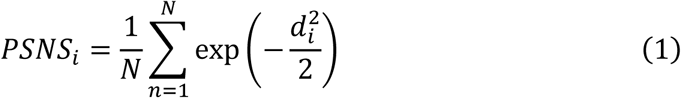

where *N* is the number of aligned homologous structures in the MSTA, and 𝑑_𝑖_ is the Cα distance between the *i*-th residue of a homologous structure and its corresponding residue in the reference structure.

#### 2.2 Evolution-guided diffusion models for pathway generation

In the second module, evolution-guided diffusion models are proposed to simulate protein folding pathway using the framework of the score-based Stochastic Differential Equations (SDE)^47, 48^. For clarity, we introduce the main concept of the pathway generation briefly in this Section. More detail about the evolution-guided forward SDE, reverse SDE, and score function is presented in the next Section.

This module consists of two stages. The first stage (**Figure 1A** bottom left) generates an initial folding trajectory using an evolution-guided conditional diffusion model. It begins with pure Gaussian noise at 𝑡 = 1 and performs 300 reverse denoising steps toward the native state at 𝑡 = 0. This results in intermediate states that are partially structured yet still contain noise. The detailed sampling procedure for stage 1 is described in **Supplementary Text S5**.

In stage 2 (**Figure 1A**, bottom right), the intermediate conformations generated in stage 1 are further refined using a fused conditional and unconditional diffusion model. To incorporate trajectory information into the refinement process, we assign dynamic per-residue weights to each intermediate conformation by measuring its structural deviation from the final stage-1 structure. The weight 𝑤_𝑖_for the *i*-th residue in a given conformation is computed using its neighbors. Specifically, if there exists a sliding window [𝑚, 𝑛] of at least three consecutive residues, i.e., for 1 ≤ 𝑚 ≤ 𝑖 ≤ 𝑛 ≤ *N*,

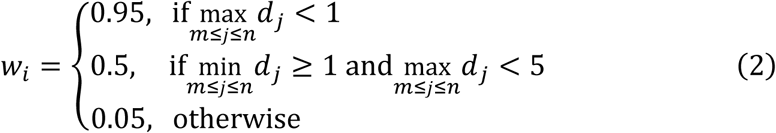

where *N* is the number of residues in the protein, and 𝑑_𝑗_ denotes the Euclidean distance between the C𝛼 atom of residue *j* in the conformation and the final stage-1 structure. This ensures that high weight is given preferentially to locally correct structural segments while discouraging isolated or short-range agreements. The resulting per-residue weights are then used to construct the adaptive dual-score function for the reverse SDE during denoising (see Section 2.5).

Reverse denoising is performed over 100 steps with a uniform noise schedule, ultimately producing a refined conformation in which well-predicted structured regions exhibit high stability, while intrinsically unstructured regions retain appropriate flexibility. More technical details of stage 2 are provided in **Supplementary Text S6**.

#### 2.3 Evolution-guided forward diffusion

##### Frame representation of protein structure

To facilitate the diffusion process, we convert the protein structure into frames. Specifically, for a protein with *N* residues, the backbone atoms (N, Cα, C) are transformed into a sequence of frames using the Gram-Schmidt process:

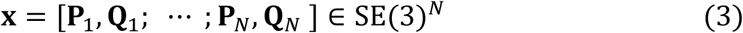

where 𝐏_𝑖_ ∈ ℝ^3^ denotes the translation vector and 𝐐_𝑖_ ∈ SO(3) is the rotation matrix of the *i*-th residue.

##### Evolution-guided forward diffusion time

The forward noising process gradually injects noise into the native protein structure, transforming it into a near-isotropic random distribution. We employ a frame-based diffusion process on the manifold SE(3)^𝑁^, following the FrameDiff^49^ method. Unlike standard homogeneous diffusion, which applies uniform noise schedules across all residues, our method modulates the noising process by incorporating evolutionary priors. This modulation is achieved through a position-specific diffusion time derived from PSNS, which assigns a distinct effective diffusion time to each residue and thereby controls the rate of noise injection on a per-residue basis. For the sequence-conditional diffusion model, the effective time step for residue *i* is calculated as follows:

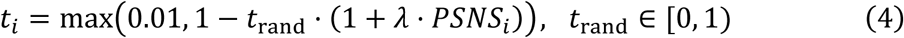

where 𝑡_rand_ is a randomly sampled scalar, and 𝜆>0 is a hyperparameter that controls the strength of the PSNS modulation. This ensures that residues with larger/smaller PSNS values are assigned smaller/larger time, resulting in weak/strong perturbations during the forward noising process.

For the unconditional diffusion model, the PSNS-based modulation is omitted. Instead, a global diffusion time *t* is uniformly applied to all residues:

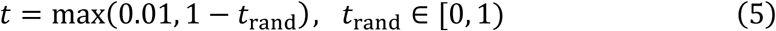

##### Forward SDE from data to noise

Based on the residue-specific diffusion time *t* defined above, we model the noise injection using a forward SDE. For the translational components, the forward SDE is given by:

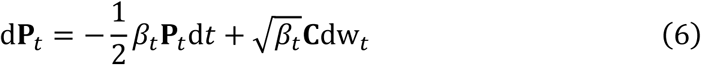

where **C** is a projection operator that eliminates global rigid-body translation of the protein; w_𝑡_ ∈ ℝ^3𝑁^ is a standard Wiener process that introduces Gaussian noise into the translational subspace; and 𝛽_𝑡_ = 𝛽_min_ + 𝑡(𝛽_max_ − 𝛽_min_). We set 𝛽_min_ = 0.1 and 𝛽_max_ = 20 to control the noise intensity. Each residue *i* is assigned a unique diffusion time 𝑡_𝑖_, which regulates the injection of noise. As 𝑡_𝑖_ increases, the structure of the corresponding backbone is gradually disrupted toward a near-isotropic noise distribution.

For the rotational components, the forward SDE is defined as:

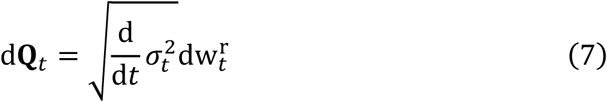

where 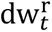 denotes the Brownian motion on the rotation group SO(3), 𝜎_𝑡_ = log(𝑡𝑒^𝜎max^ + (1 − 𝑡)𝑒^𝜎min^) with 𝜎_min_ = 0.1 and 𝜎_max_ = 1.5.

#### 2.4 Evolution-guided reverse diffusion

##### Evolution-guided reverse diffusion time

The inverse diffusion process in stage 1 begins with the random noise at 𝑡 = 1, and progressively recovers the protein backbone structure by estimating and removing the added noise. Analogous to the forward noising process, the generative (denoising) process is modulated by evolutionary priors. These priors are incorporated via position-specific time increments, Δ𝑡_𝑖_, for residue *i*:

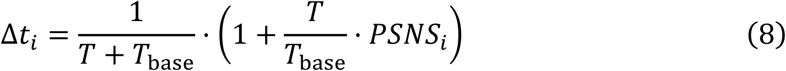

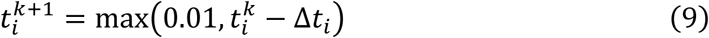

where *k* represents the current reverse diffusion step, *T* is the total number of diffusion steps, and 𝑇_base_is the base step parameter. Notably, 𝑇_base_ is the minimal number of diffusion steps required for the model to generate valid structures. In this work, 𝑇_base_is set to 100. Consistent with the training stage, this strategy ensures that residues with higher PSNS values are assigned smaller diffusion times.

For stage 2, the diffusion time is defined as a global variable *t*, uniformly applied to all residues:

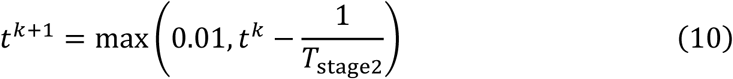

where 𝑇_stage2_ is the total number of steps in the second stage. The hyperparameters are detailed in **Supplementary Table S2**.

##### Reverse SDE from noise to data

Using the adaptive time steps derived above, the reverse SDE for the translational component **P** is formulated as:

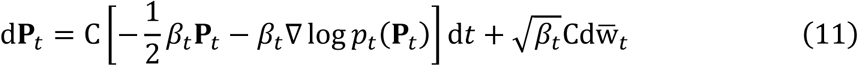

Where 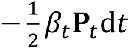 reverses the deterministic drift in the forward process for the translational component; ∇ log 𝑝_𝑡_(𝐏_𝑡_) d𝑡 is the denoising term (score function), guiding the translation toward the native structure; and 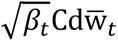 is the reverse Wiener process term, introducing necessary randomness into the denoising process.

For the rotational component **Q**, the SDE for reverse denoising is:

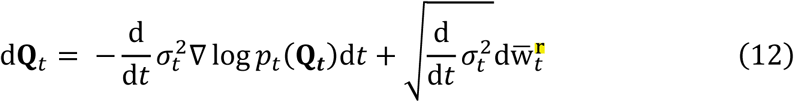

where the first term is the denoising term (driven by the score function) adapted to the derivative of the noise schedule in the rotational space, and the second term is the reverse rotational Wiener process term, introducing noise that conforms to the manifold properties of the rotation group.

To solve the reverse SDEs, we need to estimate the score function ∇ log 𝑝_𝑡_(𝐱_𝑡_). The operator ∇ denotes the gradient with respect to the noisy state variable at time *t* (i.e., 𝐏_𝑡_or 𝐐_𝑡_), and will be used hereafter unless otherwise specified. During training, we utilize the native structure 𝐱_𝟎_ = [𝐏_0_, 𝐐_0_] to compute the theoretical ground truth. According to FrameDiff^49^, the theoretical scores for the translational and rotational components **P** and **Q** can be analytically calculated as labels:

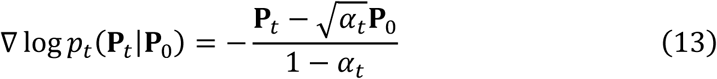

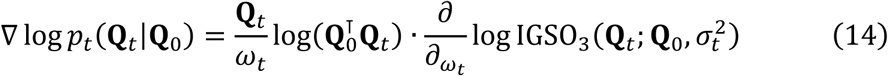

where 𝜔_𝑡_ denotes the geodesic distance on the SO(3) manifold, which is calculated via the trace of the relative rotation matrix:

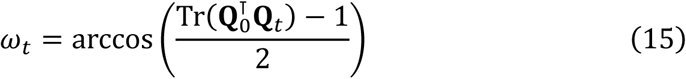

The term IGSO_3_ represents the probability density function of the Isotropic Gaussian distribution on SO(3). It is defined as an infinite series expansion based on the rotation angle 𝜔_𝑡_:

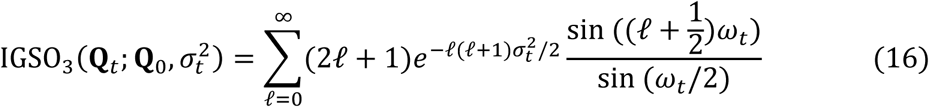

In our implementation, the infinite series is approximated by summing the first 1000 terms.

During diffusion model training, we start with a native structure 𝐱_0_, gradually add noise to obtain 𝐱_𝑡_, and then train the neural network to predict the theoretical score ∇ log 𝑝_𝑡_(𝐱_𝑡_|𝐱_0_) (defined by Eq. 13 and Eq. 14). The network’s outputs are optimized to match this score, enabling the recovery of protein structure from noise during inference. Details of the single-step reverse diffusion process are described in **Supplementary Text S4**.

#### 2.5 Stage-specific denoising strategies

During inference, since the native structure 𝐱_𝟎_ is unknown, we employ the trained networks to approximate the score functions. The sampling strategy differs between the two stages to address specific generation goals.

In stage 1, the reverse process is driven solely by the sequence-conditional score function to efficiently generate an initial folding trajectory toward the native state constraints:

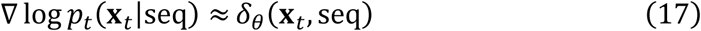

In stage 2, to balance the accurate folding of native structures with the exploration of conformational landscapes, we employ an adaptive dual-score fusion guidance mechanism (**Figure 1B**). Grounded in the principle of classifier-free guidance^47^, this formulation provides a way to estimate the conditional score by leveraging a Bayesian combination of conditional and unconditional diffusion models. The score function is defined as a linear combination of the conditional and unconditional estimators:

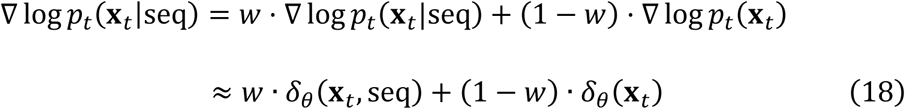

where 𝐱_𝑡_ = [𝐏_𝑡_, 𝐐_𝑡_], and the hyperparameter 𝑤 (defined in Section 2.2) regulates the balance between sequence-specific structural constraints and intrinsic disorder: near-native structures are generated by the conditional estimator when 𝑤 → 1; disordered states are generated by the unconditional estimator when 𝑤 → 0; and values in between produce conformations that have both near-native and disordered structural features, which is crucial for exploring folding intermediates.

#### 2.6 Model implementation and training

PathDiffusion trains two complementary diffusion models: a sequence-conditional model and an unconditional model. The sequence-conditional diffusion model is trained on a clustered version of the PDB database, enabling it to generate structured conformations guided by the input amino acid sequence. In contrast, the unconditional model is trained on conformational ensembles of human intrinsically disordered regions generated through coarse-grained molecular dynamics simulations from CALVADOS^29^. This specialized training allows the unconditional model to capture the characteristic flexibility and dynamical behavior of disordered segments.

##### Input representations

For the sequence-conditional model, the input representations are designed to integrate evolutionary and structural signals (**Figure 1D**). The single representation is constructed by concatenating amino acid sequence embeddings with PSNS-based diffusion time embeddings, and ESMFold-derived node embeddings. Pairwise representations are constructed by concatenating sequence embeddings, PSNS-based diffusion time embeddings, inter-residue distances computed from the noisy frame, and ESMFold-derived edge embeddings.

For the unconditional model, the input features are adjusted to avoid biasing the model towards precomputed structural priors. The single representation is constructed by concatenating a masked residue position encoding (hiding amino acid identity) with a global time embedding, where the PSNS is replaced with a uniform noise schedule. Similarly, pairwise representations are constructed by concatenating residue position encodings, a global time embedding, and inter-residue distances.

##### Structure Module

Following the ConfDiff^48^ framework, our structure module adapts the AlphaFold2^50^ architecture to a frame-based system. It integrates invariant point attention (IPA) layers, transformer layers, and multilayer perceptron (MLP) to generate and refine three-dimensional conformations. The network architecture is shown in **Supplementary Figure S1**.

The IPA module comprises 6 layers, with a single representation dimensionality of 256, a pair representation dimensionality of 128, and a hidden dimensionality of 256. Each IPA layer employs 6 attention heads, 8 query points, and 12 value points. The Transformer module contains 2 layers, each with 8 attention heads. Finally, multi-head attention and MLP layers are used for edge feature transitions and torsion angle updates, enabling the accurate prediction of folded protein conformations.

##### Loss function

The loss function is defined as:

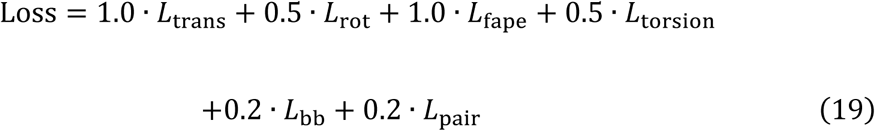

where the denoising score matching losses for translations and rotations are given by:

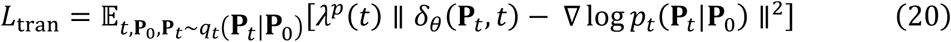

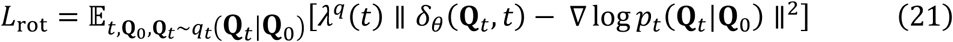

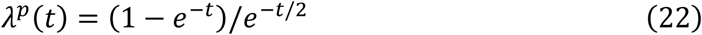

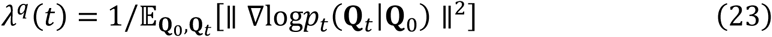

Here, ∇ log 𝑝_𝑡_(𝐏_𝑡_|𝐏_0_) and ∇ log 𝑝_𝑡_(𝐐_𝑡_|𝐐_0_) represent the theoretical ground-truth score functions for the translational and rotational components, respectively (see Eq. 13 and Eq. 14). The weighting functions 𝜆^𝑝^(𝑡) and 𝜆^𝑞^(𝑡) are employed to balance the loss magnitudes across different diffusion time steps, ensuring stable optimization.

These losses are combined with auxiliary structural losses: a frame aligned point error (FAPE) loss (𝐿_fape_) for global fold correctness, a torsion angle loss (𝐿_torsion_) for protein backbone dihedrals (φ, ψ, ω), a backbone coordinate MSE loss (𝐿_bb_), and a pairwise distance loss (𝐿_pair_) for local atomic interactions. This multi-component loss ensures the model learns accurate denoising dynamics while generating physically plausible protein structures.

## Data availability

The benchmark dataset can be downloaded at https://yanglab.qd.sdu.edu.cn/PathDiffusion/

## Software availability

The source code and model parameters are available at https://github.com/YangLab-SDU/PathDiffusion

## Acknowledgements

This work is supported by the following funding sources: National Natural Science Foundation of China (NSFC T2225007, 62503282, 32430063, 62501364), Postdoctoral Fellowship Program and China Postdoctoral Science Foundation (2025M773121, BX20240212, 2025M783122), the Basic Research Program of Jiangsu Province (BK20250432), Qingdao Postdoctoral Applied Research Project, and Fundamental Research Funds for the Central Universities.

## Supplementary Information

### Text S1. Dataset

#### 1.1 Training dataset

The PathDiffusion network consists of two models: sequence-conditional diffusion model and conditional diffusion model, which are used to fold structured regions and unstructured regions, respectively. Two training datasets were constructed accordingly to train both models.

##### Training dataset of structured proteins

The training and validation datasets were constructed using single-chain structures from the Protein Data Bank (PDB) to train the sequence-conditional diffusion model. We used entries released prior to April 10, 2024, for training, and those deposited between April 10, 2024, and April 16, 2025, for validation. Following the data processing strategy proposed in ConfDiff^1^, we expanded our dataset by extracting and preserving all available structural models within each PDB entry, which allows us to fully leverage the conformational ensembles provided by experimental techniques such as solution NMR. To ensure data quality, we applied the following filtering criteria: (1) sequence length is 20-500 residues; (2) chains with >30% missing backbone atoms were discarded; and (3) chains with random coil ratio >50% were removed. After strictly removing proteins overlapping with the benchmark test set, we obtained a final dataset of 538,380 structures for training and 40,835 for validation. All chains were clustered at a 70% sequence identity threshold. In each training iteration, the data loader first randomly selected a cluster and then extracted a specific structural conformation from that cluster.

##### Training dataset of unstructured proteins

We used intrinsically disordered regions (IDRs) from CALVADOS simulations to train the unconditional diffusion model^2^. These simulations covered nearly all intrinsically disordered regions in the human proteome, but only contained Cα atoms. We selected protein chains ranging from 20 to 500 in length and used pdbfixer^3^ to restore the coarse-grained structures into all-atom models. The final dataset was split into a training set and a validation set in a 9:1 ratio: 24,206 structures were used for training, and 2,688 structures were used for validation.

#### 1.2 Test dataset

Four benchmark datasets were constructed to evaluate PathDiffusion: FP52, MD12, LT20 and IDP50.

##### FP52

The original FoldPAthreader benchmark contains 30 proteins. We expanded this dataset to include 22 more proteins, resulting in 52 proteins (denoted by FP52). The folding information for these proteins were collected from the literature, derived from experimental methods including hydrogen-deuterium exchange mass spectrometry (HX-MS), circular dichroism (CD), fluorescence resonance energy transfer (FRET), nuclear magnetic resonance (NMR), and related approaches. The sequence lengths of these 52 proteins range from 43 to 363 residues. A detailed list of proteins and their annotations are provided in **Supplementary Table 1**.

##### MD12

The dataset includes 12 small proteins, ranging from 10 to 80 residues in length (denoted by MD12). Their structures have been extensively simulated by MD on the Anton supercomputer^4^. Each protein was simulated in 1 to 4 independent runs, with a total simulation time of ∼8 milliseconds. This extended sampling captured more than 400 folding and unfolding events, yielding sufficient statistics to characterize folding dynamics. The structural data are represented at the Cα-atom level, which provides a coarse but informative description of the folding process.

##### LT20

This dataset comprises 20 proteins known to exhibit local folding-unfolding transition (denoted by LT20), from the BioEmu study^5^. In each instance, a specific segment of at least 8 residues is defined as undergoing a transition from a folded to an unfolded state.

##### IDP50

This dataset contains 50 intrinsically disordered proteins from PDB (denoted by IDP50). The selection criteria included: (1) more than 70% of the residues are random coils; (2) sequence length is from 50 to 100; and (3) pairwise sequence identity is lower than 70%. This screening process yielded 681 candidate proteins, from which 50 were randomly selected.

### Text S2. Folding pathway evaluation metrics

To quantitatively assess the folding sequence, we calculated the experimentally defined early folded region (EFR) and late folded region (LFR) lDDT scores by comparing sampled conformations with the native. Specifically, the EFR lDDT was computed exclusively based on the intra-regional contacts within the early folded residues. In contrast, the calculation of the LFR lDDT was designed to cover the entire late-stage folding process; therefore, it considered not only the intra-regional contacts within the late folded regions but also the inter-regional contacts formed between the early and late regions. This ensures that the LFR lDDT reflects both the local structure of the late regions and their packing with the already folded core. A conformation with an lDDT of the EFR being more than 10% higher than that of the LFR was defined as a true positive conformation (TPC).

### Text S3. Z-score analysis of folding order

To quantitatively validate the predicted folding pathways against experimental evidence, we performed a comparative Z-score analysis. The experimental data were derived from HX-MS datasets reported in the literature, utilizing specific metrics appropriate for each protein’s experimental design:

- **sIGPS (PDB: 2C3Z):** Data were obtained from equilibrium pulse-labeling experiments in urea^6^. The metric used was the percentage of protected amide protons within the stable equilibrium intermediate populated at ∼5 M urea. Higher protection levels indicate regions that form the stable structural core.
- 𝜶**TS (PDB: 1BKS):** Data were derived from kinetic refolding pulse-labeling experiments^7^. The metric used was the relative population of protected species for each peptide segment, measured after 2 seconds of refolding. A higher population indicates regions that fold rapidly into the kinetic intermediate.
- **Aldolase (PDB: 1ADO):** Data were collected during refolding via dialysis^8^. The metric used was the inverse of the dialysis time (1/𝑡_1/2_) required for a segment to achieve protection. Since earlier protection occurs at higher denaturant concentrations (shorter dialysis times), the inverse time serves as a proxy for folding priority.

For the PathDiffusion predictions, we quantified the folding timing for each residue by analyzing the generated trajectories. All sample conformations were globally aligned to the final state. We calculated a folding proportion for each residue, defined as the percentage of conformations in the trajectory where the C𝛼 atomic distance to the corresponding residue of final state was less than 3 Å.

To ensure direct comparability with the low-resolution MS data, the residue-level folding proportions from PathDiffusion were averaged over the specific segments defined in the respective experimental studies. Finally, both the segment-averaged experimental values and the segment-averaged predicted values were standardized into Z-scores (𝑍 = (𝑥 − 𝜇)/𝜎). In the comparative plots, horizontal lines represent the Z-scores for experimentally resolved peptide segments, while diagonal lines connect these segments, indicating regions where experimental data were unavailable. A positive Z-score signifies an early folded region, while a negative Z-score signifies a late folded region.

### Text S4. Algorithm 1: reverse diffusion

This algorithm outlines the single-step reverse denoising operation, which implements the numerical discretization of the reverse-time SDEs defined in the reverse denoising process in Section 2.4. It takes the current conformational state 𝑥^(^*^𝑡^*^)^, the predicted score 𝑠 (derived from the neural network output), and the current time step 𝑡 as inputs. For each residue 𝑙, the algorithm updates translational coordinates in Euclidean space and rotational orientations on the *SO*(3) manifold separately, incorporating the residue-wise diffusion time to transition the structure from state 𝑥^(𝑡)^ to 𝑥^(𝑡−𝛥𝑡)^.

#### Algorithm 1

PathDiffusion Reverse Diffusion Process

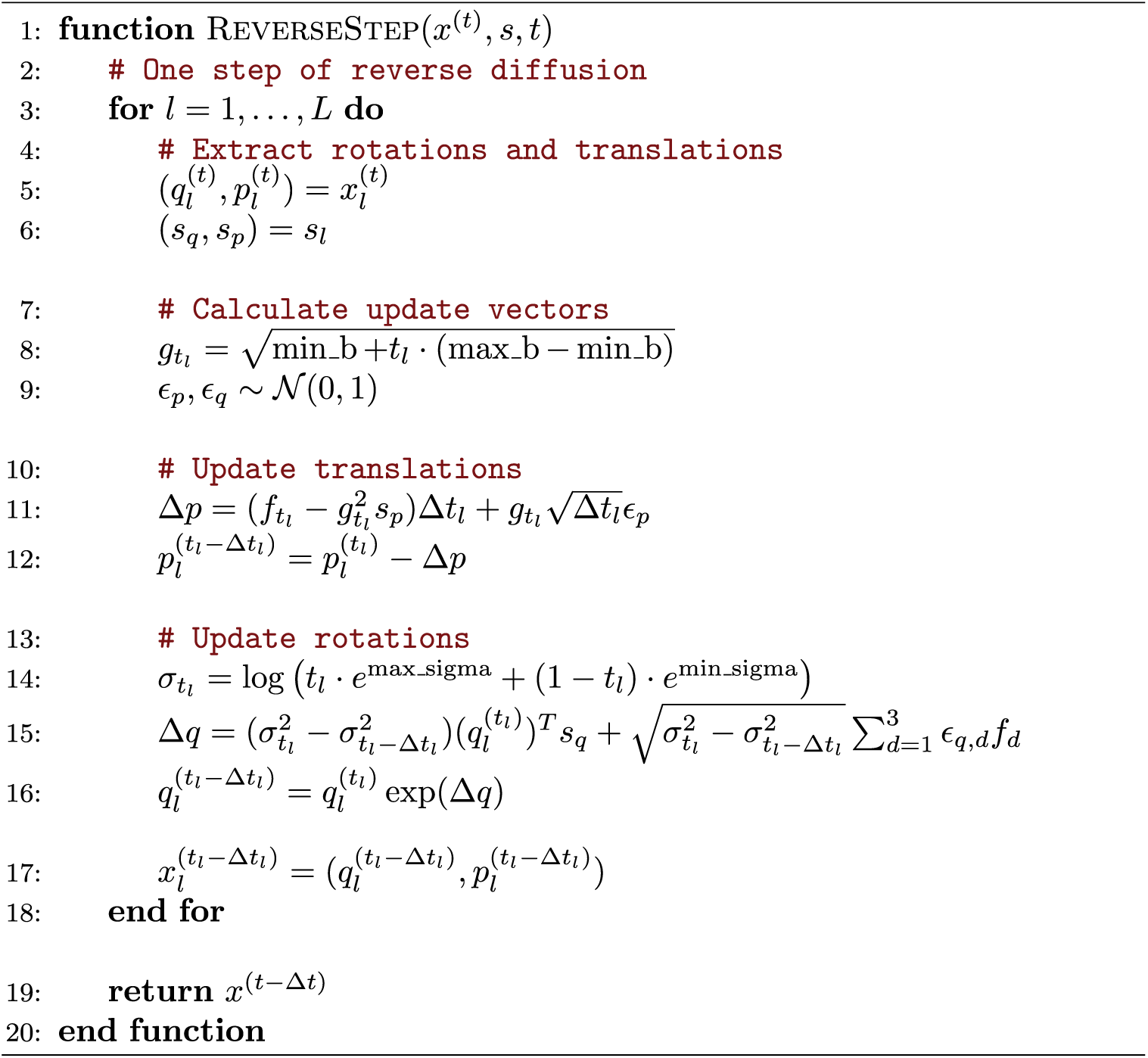

### Text S5. Algorithm 2: sampling process in stage 1

This algorithm outlines the sampling process in stage 1. The *SampleStage1* initializes the protein backbone with random noise for both rotations and translations. It then defines a position-specific diffusion time based on the input position-specific noise schedules (PSNS). In each iteration 𝑘, the model predicts the denoised structure 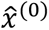, computes the conditional score 𝑠^cond^, and updates the conformation using the *ReverseStep* function. Intermediate structures are saved when the process enters the designated time window to record the folding trajectory.

#### Algorithm 2

PathDiffusion Stagel Generation

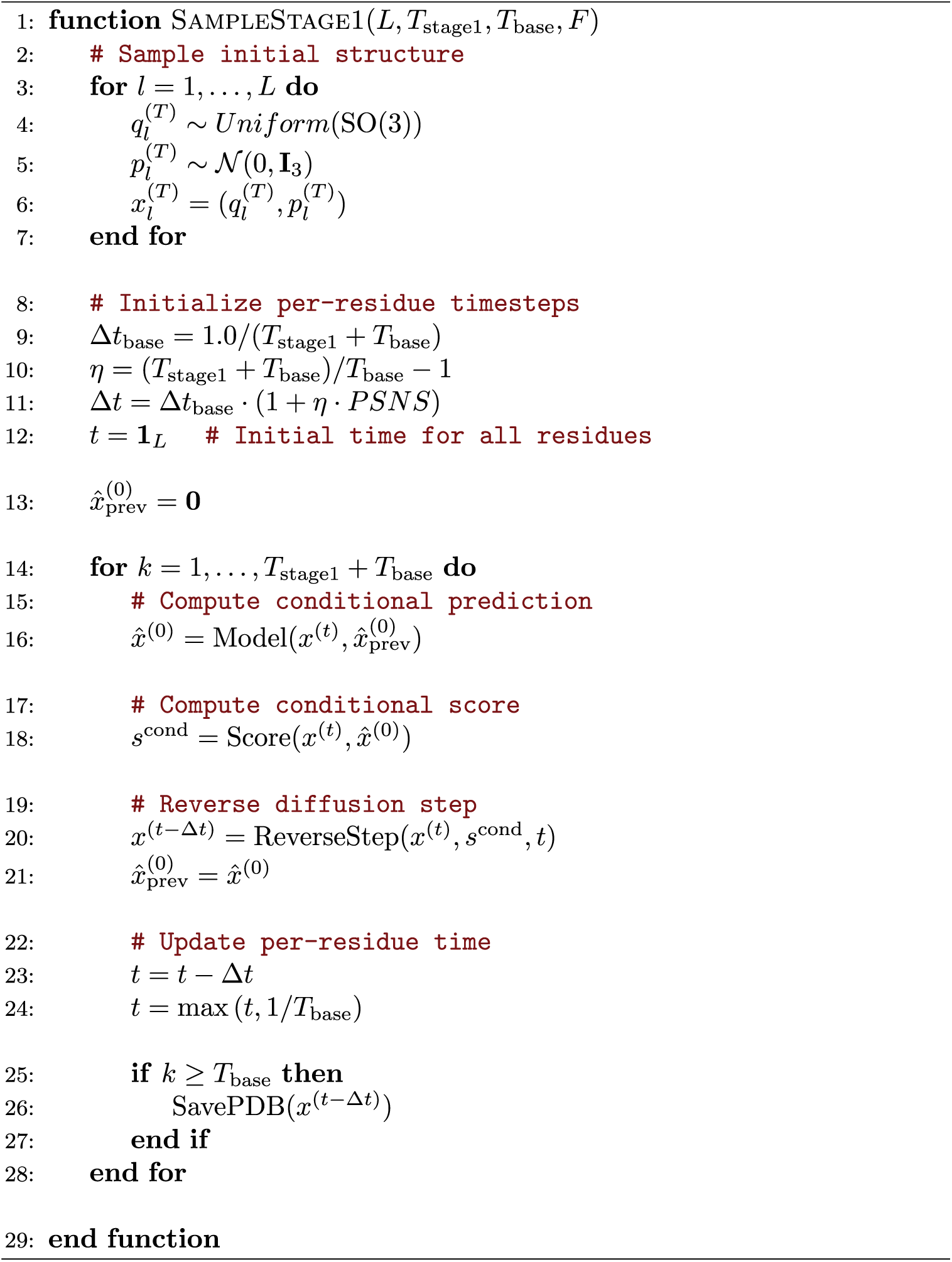

### Text S6. Algorithm 3: sampling process in stage 2

This algorithm presents the adaptive dual-score fusion strategy employed in the second stage to refine the trajectory dynamics. The algorithm first computes dynamic weights 𝜔 for each residue based on the structural consistency with the predicted final state of stage1 (Lines 2-15). These weights govern the interpolation between the sequence-conditional score 𝑠^cond^ and the unconditional score 𝑠^unc^ (Line 28).

#### Algorithm 3

PathDiffusion Stage2 Generation

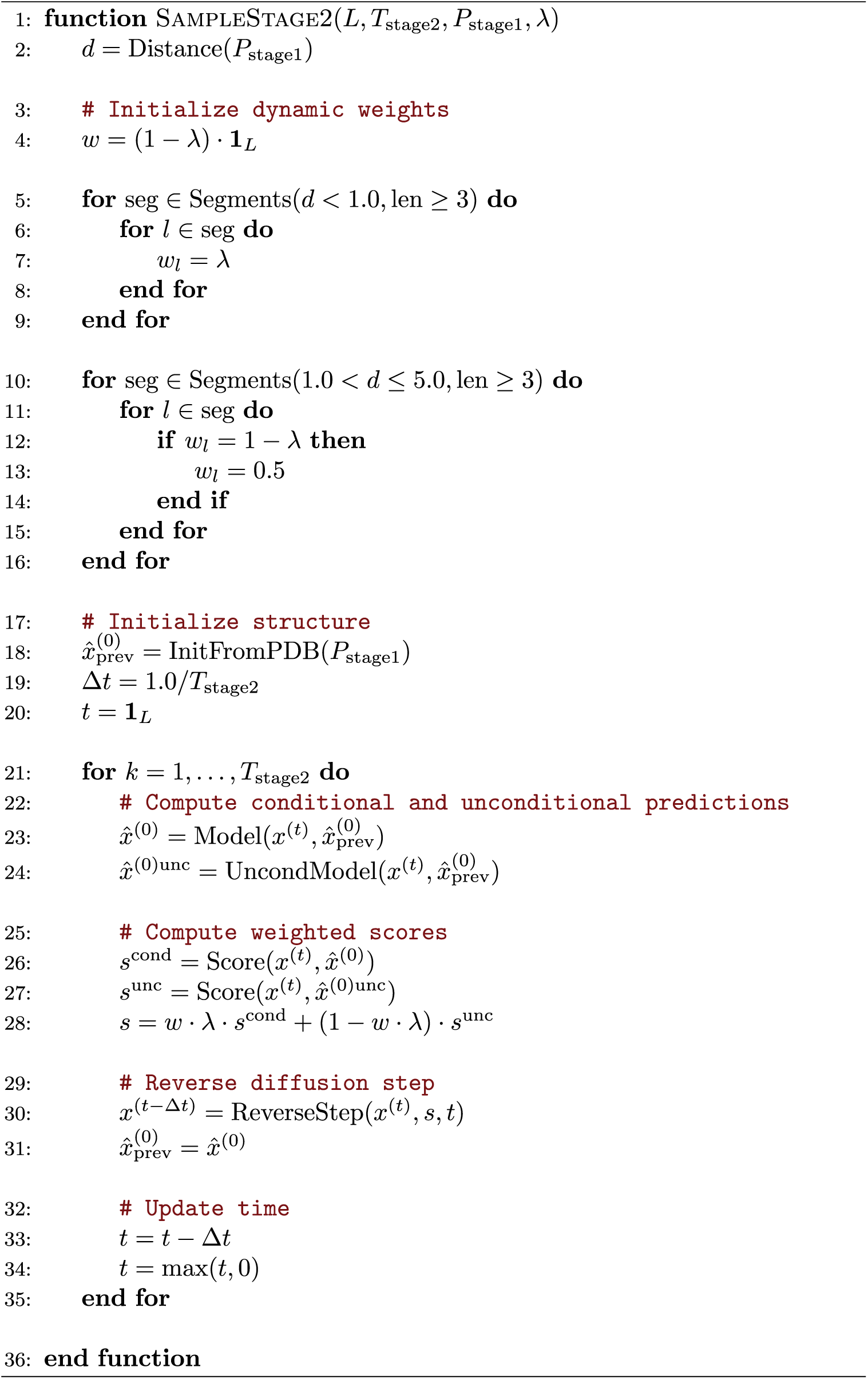

**Table S1.** Detailed information of FP52 dataset. Early folded residues were annotated based on evidence from the literature.

| Protein name | PDB ID | Length | Early fold residues | Reference |
| --- | --- | --- | --- | --- |
| **Barstar** | 1AB7_A | 89 | 1-64 | 10.1073/pnas.94.3.826<br>10.1006/jmbi.1996.0359 |
| **Muscle-type aldolase** | 1ADO_A | 363 | 1-57, 131-298 | 10.1021/bi027388q<br>10.1006/jmbi.1999.3251 |
| **Apo-azurin** | 1AIZ_A | 129 | 27-36, 89-98,<br>106-112, 121-129 | 10.1002/prot.22099 |
| **Im7** | 1AYI_A | 86 | 12-45, 66-79 | 10.1038/nsb757<br>10.1016/j.jmb.2004.01.004<br>10.1016/j.jmb.2009.07.085 |
| **PDZ-3 domain** | 1BE9_A | 115 | 10-92 | 10.1016/j.jmb.2004.11.040 |
| **Barnase** | 1BGS_A | 110 | 1-22, 53-75,<br>85-110 | 10.1021/bi0362267<br>10.1073/pnas.0408226102 |
| **Tryptophan synthase** | 1BKS_A | 255 | 16-61, 75-101,<br>110-127 | 10.1016/j.jmb.2005.01.064<br>10.1016/j.jmb.2004.05.062 |
| **Myotrophin** | 1D9S_A | 136 | 69-136 | 10.1073/pnas.0604653104 |
| **CTL9** | 1DIV_A | 90 | 66-90 | 10.1002/prot.22099 |
| **Ckshs1** | 1DKT_A | 72 | 1-21, 50-72 | 10.1016/s0022-2836(02)01202-0 |
| **FAS-associated death domain** | 1E3Y_A | 104 | 1-32, 47-79 | 10.1007/s00249-011-0756-6 |
| **MAK33** | 1FH5_L | 105 | 1-32, 38-48,<br>65-91, 97-105 | 10.1073/pnas.0802809105 |
| **Flavodoxin** | 1FTG_A | 168 | 1-8, 47-53, 80-87,<br>98-110, 139-143, 151-168 | 10.1006/jmbi.1998.2045 |
| **Immune globulin G** | 1HCV_A | 116 | 14-23, 32-51,<br>65-71, 77-98, 105-116 | 10.1021/bi0009082 |
| **HIV-1 ribonuclease H** | 1HRH_A | 125 | 13-21, 46-72, 89-110 | 10.1002/pro.5560071014 |
| **Cytochrome c** | 1I5T_A | 104 | 1-14, 61-69, 87-104 | 10.1073/pnas.1706196114<br>10.1073/pnas.0501043102<br>10.1110/ps.072922307 |
| **FKBP12** | 1J4H_A | 107 | 21-30, 56-65, 71-76, 96-107 | 10.1006/jmbi.1999.2942 |
| **Apomyoglobin** | 1MBC_A | 153 | 7-18, 24-44, 61-78, 102-142 | 10.1073/pnas.0804033105<br>10.1073/pnas.1105682108<br>10.1126/science.8235610 |
| **FHA Domain** | 1MZK_A | 122 | 4-11, 52-70, 97-119 | 10.1016/j.jmb.2006.08.090 |
| **Acyl-CoA binding protein** | 1NTI_A | 86 | 1-36, 65-86 | 10.1073/pnas.0509100103<br>10.1016/j.jmb.2013.10.031 |
| **Onconase** | 1ONC_A | 103 | 17-36, 55-99 | 10.1021/bi961085c |
| **Neurogenic locus Notch protein** | 1OT8_A | 209 | 25-150 | 10.1002/pro.9<br>10.1016/j.str.2006.06.013 |
| **BPTI** | 1QLQ_A | 58 | 18-58 | 10.1073/pnas.1503909112 |
| **TOP7** | 1QYS_A | 91 | 23-91 | 10.1016/j.bpj.2008.11.004<br>10.1073/pnas.0911844107 |
| **Ribosomal protein S6** | 1RIS_A | 97 | 1-67 | 10.1073/pnas.0401672101 |
| **Ribonuclease H** | 1RNH_A | 148 | 42-117 | 10.1126/science.1116702<br>10.1016/j.jmb.2009.05.085<br>10.1073/pnas.1305887110 |
| **Fyn SH3 domain** | 1SHF_A | 59 | 25-30, 36-41,<br>47-51 | 10.1073/pnas.0404436101 |
| **Staphylococcal nuclease** | 1STN_A | 136 | 1-36, 66-90 | 10.1016/j.bbrc.2008.08.073<br>10.1073/pnas.91.2.449 |
| **Polyubiquitin-C** | 1UBQ_A | 76 | 1-16, 22-34,<br>64-76 | 10.1073/pnas.89.6.2017<br>10.1021/bi00161a019 |
| **FF domain** | 1UZC_A | 69 | 1-51 | 10.1016/j.jmb.2007.06.012<br>10.1126/science.1191723<br>10.1016/j.jmb.2005.04.067 |
| **Rd-apocytochrome b562** | 1YYJ_A | 104 | 22-43, 65-89 | 10.1073/pnas.0501372102<br>10.1016/j.bpj.2010.01.052 |
| **Acylphosphatase** | 2ACY_A | 98 | 48-155 | 10.1016/j.jmb.2006.06.041 |
| **sIGPS** | 2C3Z_A | 220 | 1-20, 36-68,<br>76-81 | 10.1016/j.jmb.2007.02.027 |
| **CTX III** | 2CRT_A | 60 | 19-39, 49-60 | 10.1074/jbc.273.17.10181 |
| **CspB** | 2F52_A | 67 | 1-19, 46-53 | 10.1016/j.jmb.2004.04.011 |
| **DESIGNED TOP7** | 2GJH_A | 57 | 19-57 | 10.1073/pnas.0708411105 |
| **Ubq-UIM** | 2KDI_A | 114 | 11-80 | 10.1016/j.bpc.2011.05.004 |
| **T4 Lysozyme** | 2LZM_A | 164 | 1-11, 65-164 | 10.1016/j.jmb.2006.10.048 |
| **PinWW** | 2M8I_A | 43 | 27-43 | 10.1073/pnas.0905466106 |
| **Gelation factor** | 2N62_L | 221 | 1-112 | 10.1016/j.pnmrs.2016.10.002 |
| **tANK** | 2RFM_A | 183 | 87-183 | 10.1073/pnas.0710657105 |
| **Thioredoxin** | 2TRX_A | 108 | 1-90 | 10.1016/j.bbapap.2014.11.004 |
| **Beta-lactoglobulin** | 3BLG_A | 162 | 91-162 | 10.1038/84145<br>10.1006/jmbi.1999.3515 |
| **CHEY** | 3CHY_A | 128 | 1-60 | 10.1016/S1359-0278(96)00011-9 |
| **Chymotrypsin Inhibitor 2** | 3CI2_A | 64 | 12-34, 45-50 | 10.1016/j.polymer.2003.10.092<br>10.1073/pnas.0702265104 |
| **Monellin** | 3PXM_A | 96 | 10-47 | 10.1021/acs.chemrev.1c00704 |
| **RNase T1** | 3RNT_A | 104 | 56-61, 75-81,<br>86-92 | 10.1002/pro.5560060702 |
| **SasG** | 3TIP_A | 132 | 53-132 | 10.1073/pnas.1608762113 |
| **Utrophin** | 3UUL_A | 115 | 1-26, 74-104 |  |
| **Haloalkane dehalogenase** | 5UY1_A | 290 | 1-127, 212-238, 275-290 | 10.1126/sciadv.aas9098 |
| **Lysozyme C** | 6LYZ_A | 129 | 1-39, 85-129 | 10.1110/ps.8.1.35<br>10.1096/fasebj.10.1.8566530<br>10.1073/pnas.92.20.9029 |
| **Plastocyanin** | 9PCY_A | 99 | 1-6, 13-15, 18-31, 69-83, 93-99 | 10.1021/bi00097a005 |

**Table S2.** Hyperparameter of PathDiffusion.

| Category | Parameter | Value | Description |
| --- | --- | --- | --- |
| **Model Architecture** | num_ipa_blocks | 6 | Number of IPA blocks |
|  | c_s | 256 | Dimension of single representation |
|  | c_z | 128 | Dimension of pair representation |
|  | c_hidden | 256 | Hidden dimension |
|  | c_skip | 64 | Dimension of skip connections |
|  | no_heads | 6 | Number of attention heads in IPA |
|  | no_qk_points | 8 | Number of query/key points for IPA |
|  | no_v_points | 12 | Number of value points for IPA |
|  | seq_tfmr_num_layers | 2 | Number of Transformer layers |
|  | seq_tfmr_num_heads | 8 | Number of heads in Transformer |
| **Embeddings** | node_emb_size | 256 | Dimension of node embeddings |
|  | edge_emb_size | 128 | Dimension of edge embeddings |
|  | time_emb_size | 64 | Dimension of sinusoidal time embeddings |
|  | res_idx_emb_size | 64 | Dimension of residue index embeddings |
| **Inference** | $T$ | 300 | Steps for stage1 |
| | $T_{\text{base}}$ | 100 | Minimum number of diffusion steps |
| | $T_{\text{stage2}}$ | 100 | Steps for stage2 |
|  | starting_steps | 0 | Define the start and end points of the reverse diffusion trajectory, allowing flexible sampling of arbitrary partial of the trajectories |
|  | final_steps | 300 |  |

**Figure S1.**
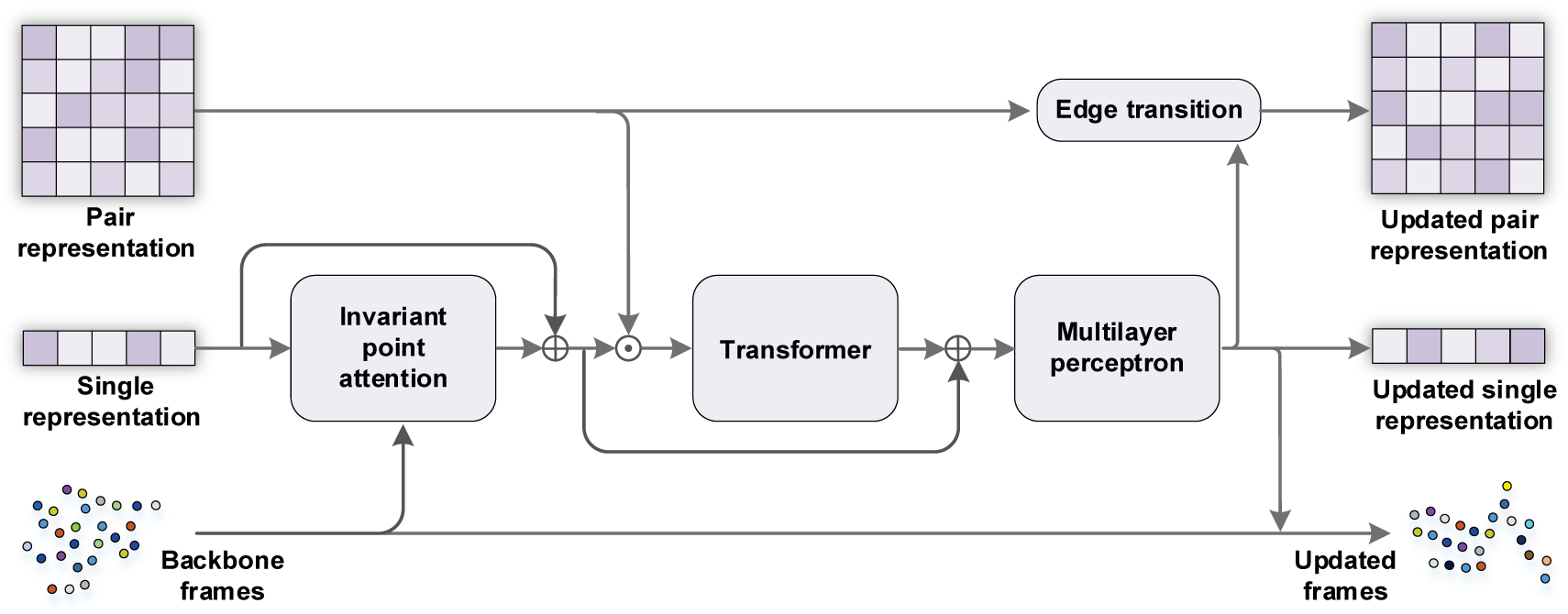
Flowchart of the structure module. The module takes the single representation, pair representation, and backbone frames as inputs. It utilizes Invariant Point Attention (IPA), Transformer blocks, and a Multilayer Perceptron (MLP) to iteratively update the representations and backbone geometry.

**Figure S2.**
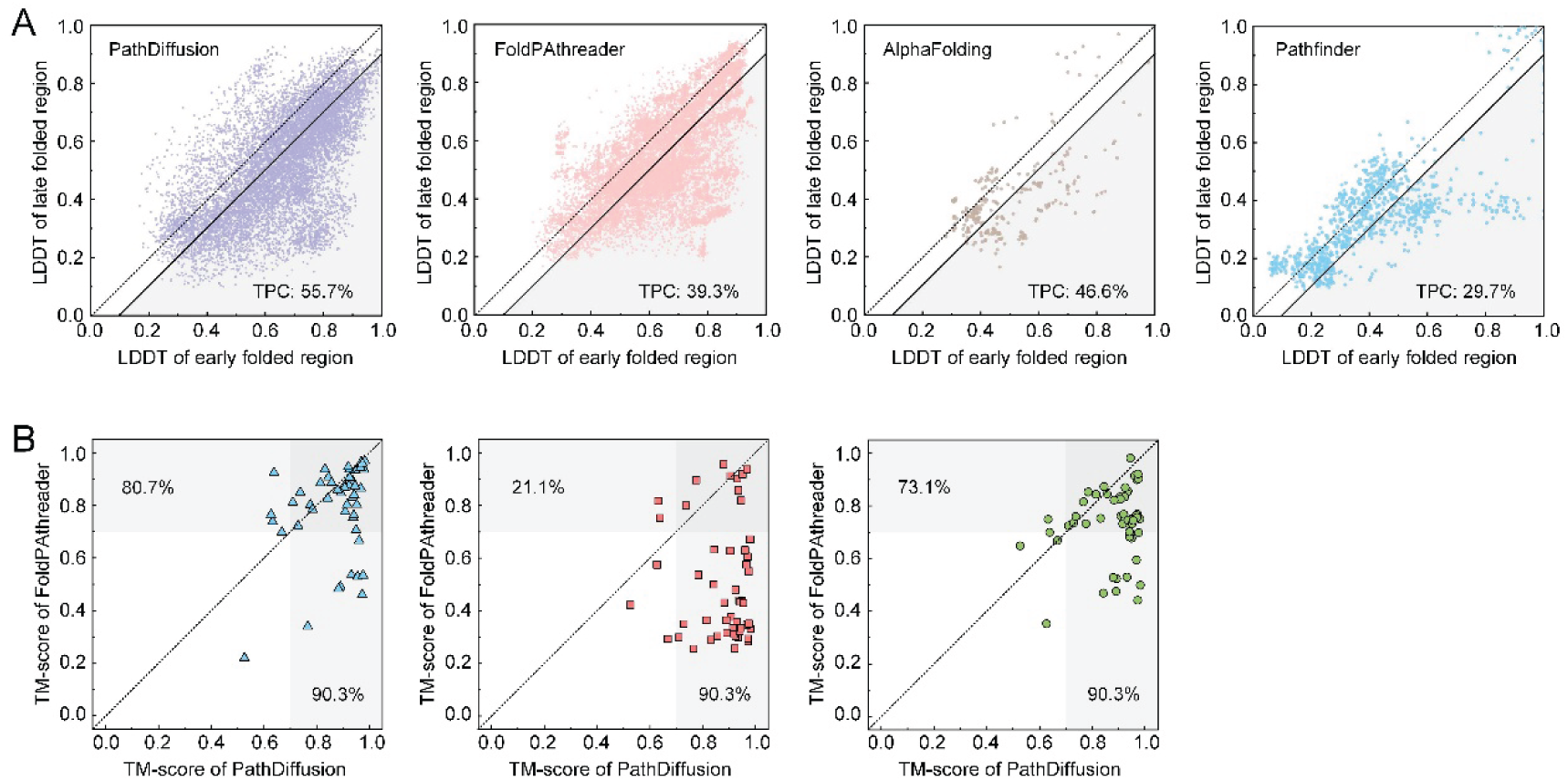
Performance evaluation of folding pathway and final state on FP52. **(A)** Scatter plots of lDDT scores for early folded regions (EFRs) versus late folded regions (LFRs). The grey shaded region represents true positive conformations (TPCs). The percentages indicate the overall proportion of TPCs for each method. **(B)** Head-to-head comparison of final state TM-scores between PathDiffusion and other methods. The grey shaded area represents the near-native (TM-score ≥ 0.7).

**Figure S3.**
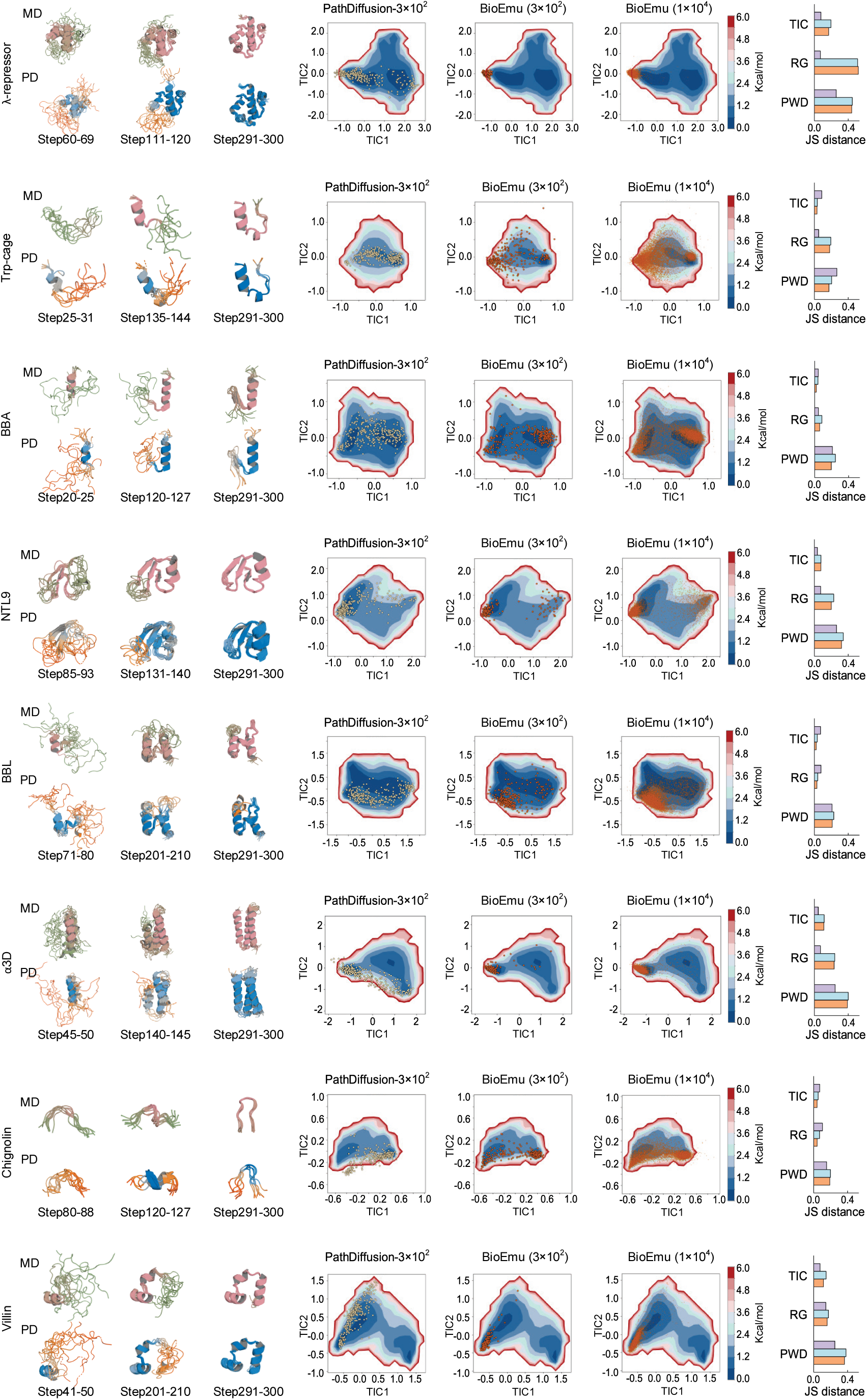
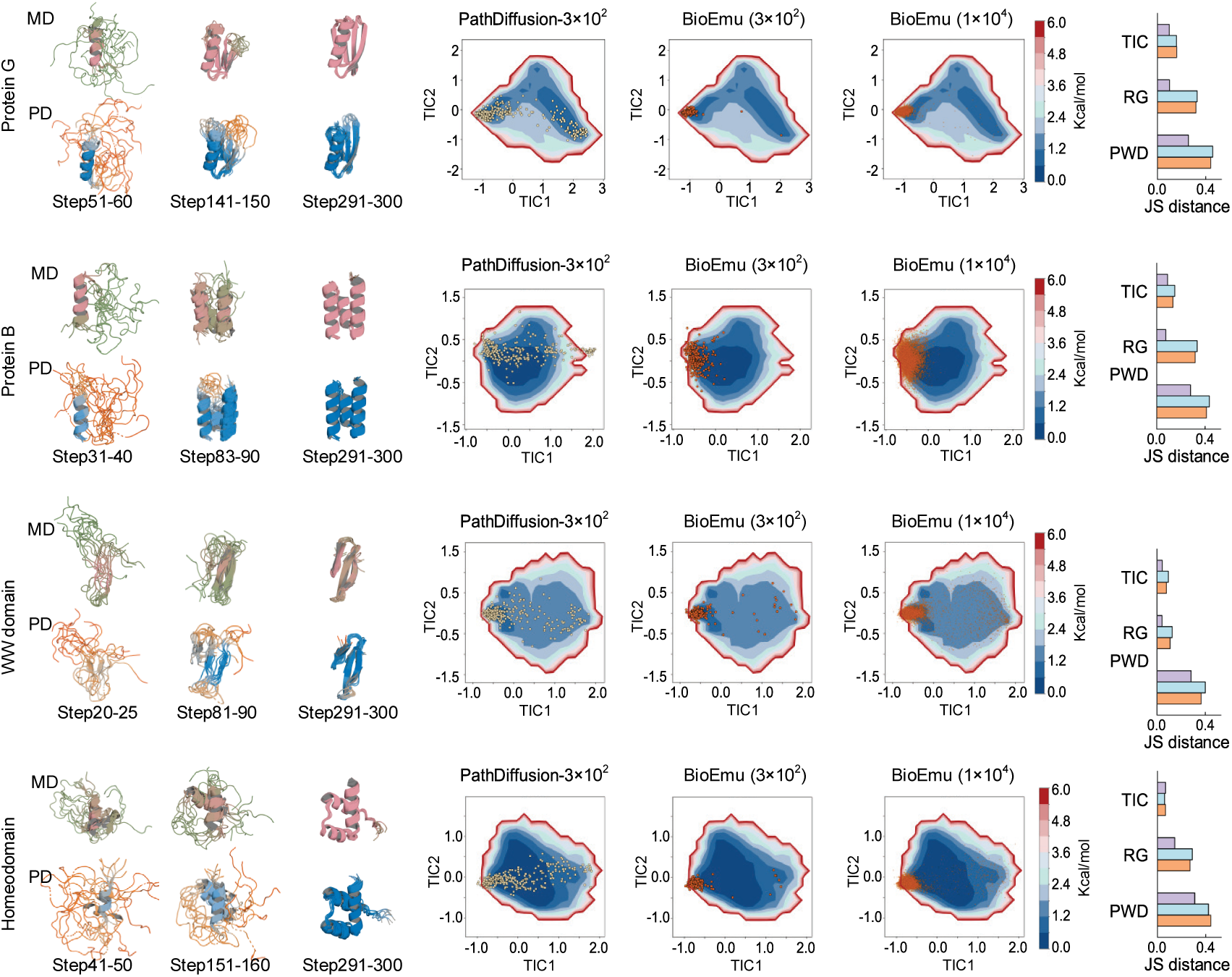
Detailed evaluation on molecular dynamics simulation dataset (MD12). Left panels: Visual comparison of MD-simulated ensembles and PathDiffusion folding trajectories at indicated sampling steps. **Middle panels:** Projection of sampled conformations (scatter points) onto the MD-derived 2D free energy surface. **Right panels:** Corresponding Jensen-Shannon distance (JSD) for each protein across the three metrics (TIC, RG, and PWD). Purple represents PathDiffusion (3×10^2^), light blue represents BioEmu (3×10^2^), and orange represents BioEmu (1×10^4^).

**Figure S4.**
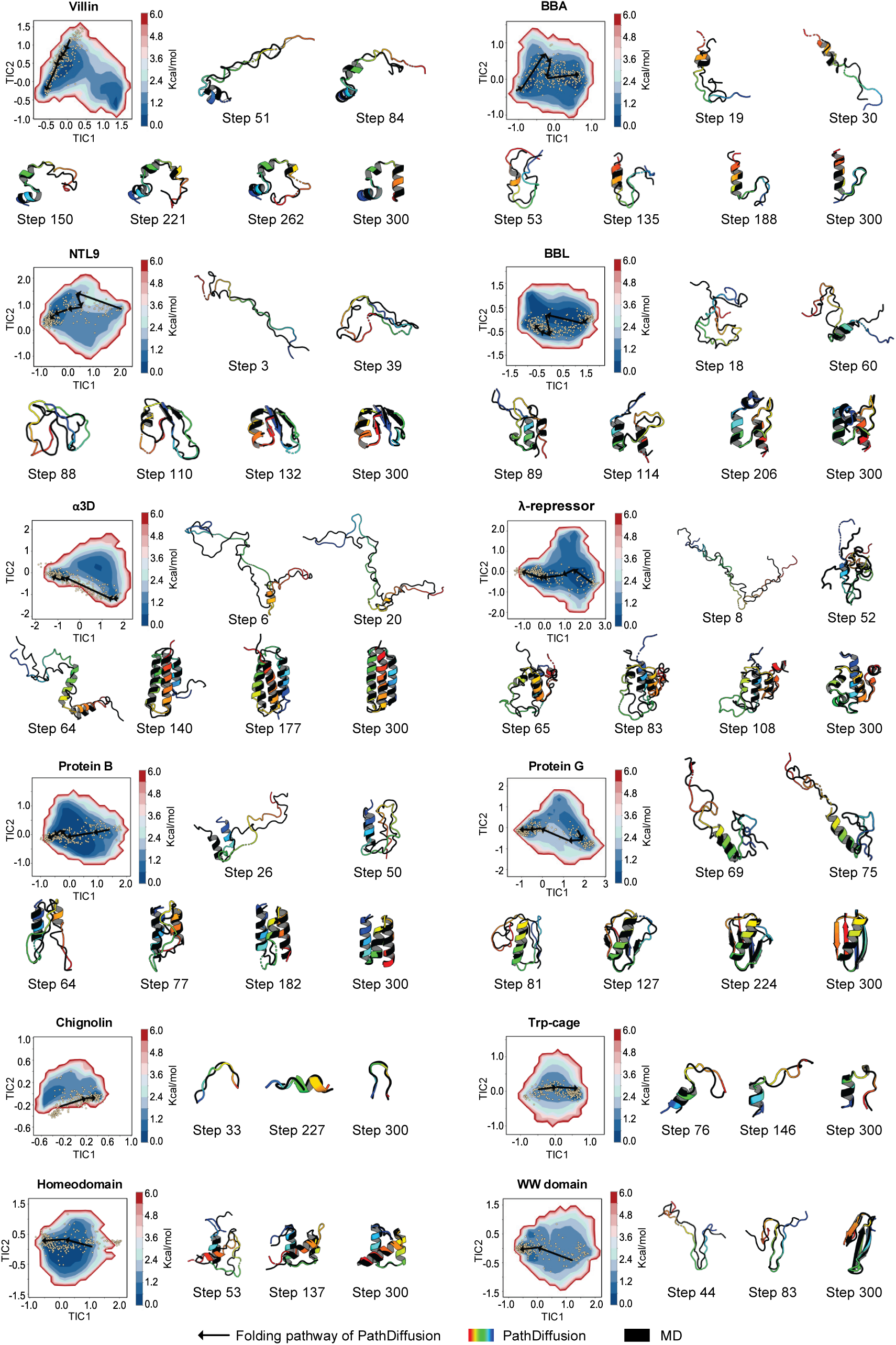
Visualization of PathDiffusion folding pathways. For each protein, the 2D free energy surface is projected onto the first two Time-lagged Independent Components (TICs) derived from ground-truth MD simulations. Yellow scatter points represent the conformations sampled by PathDiffusion, while the black arrow indicates the direction of the predicted folding trajectory. Representative snapshots at specific diffusion steps illustrate the structural evolution. In each snapshot, the PathDiffusion-generated conformation (rainbow colored) is superimposed onto the most structurally similar conformation identified in the MD simulation (black).

**Figure S5.**
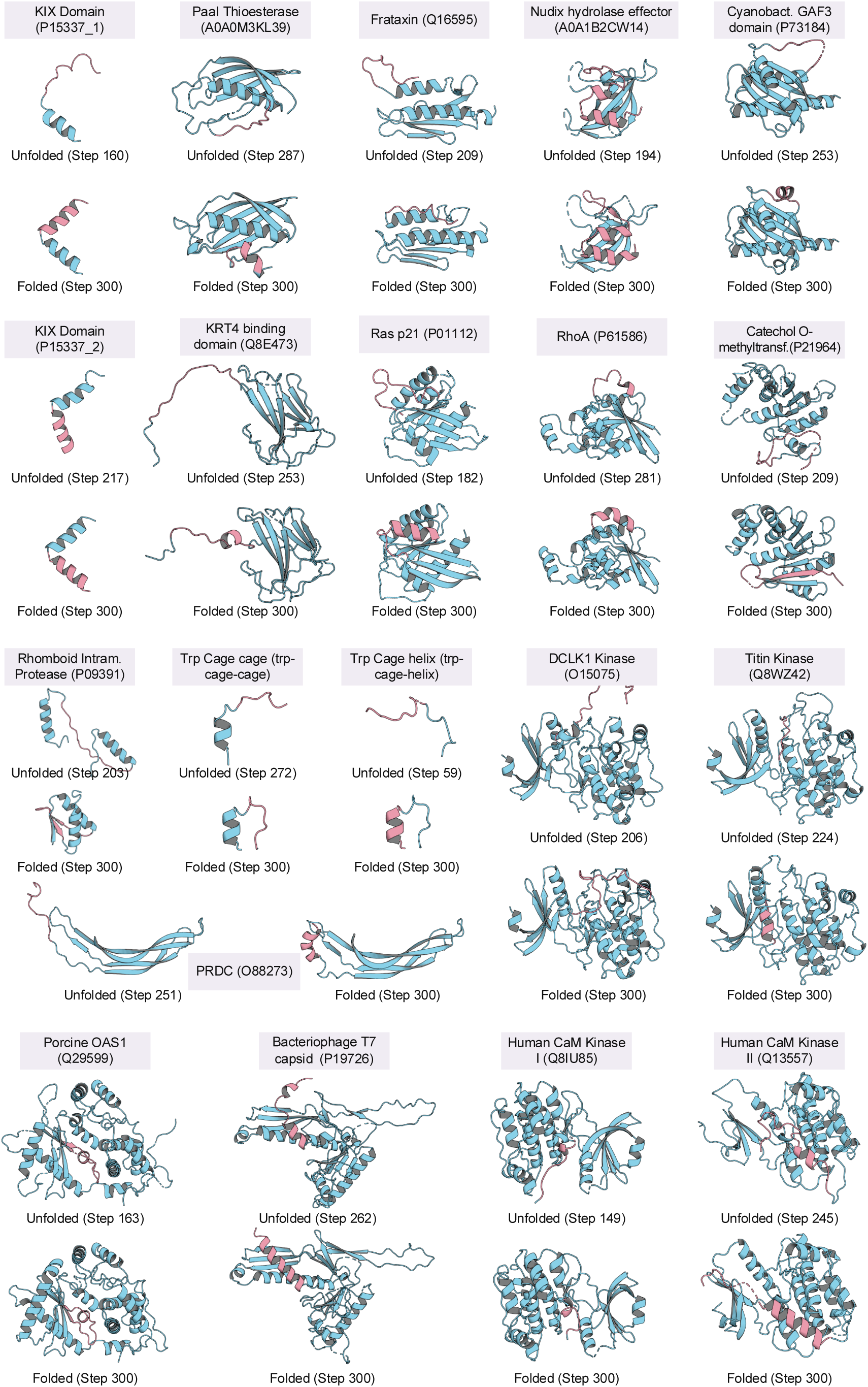
Visualization of predicted results for 20 proteins in the local folding-unfolding transition dataset (LT20). For each target, the figure displays a representative intermediate conformation (labeled “Unfolded” with the specific step number) alongside the final generated structure (labeled “Folded”, Step 300). Pink segments highlight the specific regions of the protein known to undergo local unfolding or exhibit intrinsic disorder.

**Figure S6.**
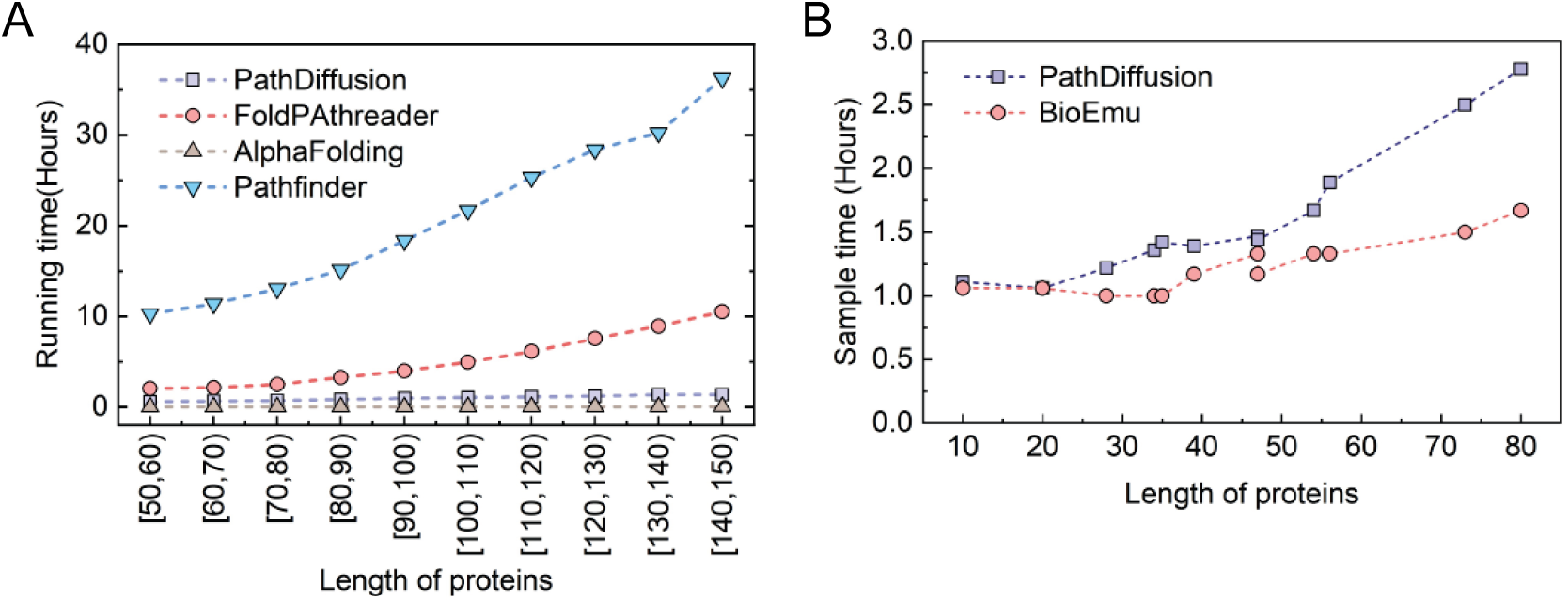
Computational efficiency analysis. **(A)** Comparison of the running time required to sample folding pathways across varying protein lengths. **(B)** Comparison of sample time for generating conformational ensembles (10,000 samples) for the 12 MD-simulated proteins.

## Notes

### Competing Interest Statement

The authors have declared no competing interest.

